# Length-limitation of astral microtubules orients cell divisions in intestinal crypts

**DOI:** 10.1101/2022.09.02.506333

**Authors:** Saleh Jad, Fardin Marc-Antoine, Frenoy Olivia, Soleilhac Matis, Gaston Cécile, Cui Hongyue, Dang Tien, Noémie Gaudin, Vincent Audrey, Minc Nicolas, Delacour Delphine

## Abstract

Planar spindle orientation is critical for epithelial tissue organization, and generally instructed from the long cell shape axis or cortical polarity domains. We introduced mouse intestinal organoid crypts to study spindle orientation in a monolayered mammalian epithelium. Although spindles were planar in this tissue, mitotic cells remained elongated along the apico-basal axis and polarity complexes were segregated to basal poles, so that spindles oriented in an unconventional manner, orthogonal to both polarity and geometric cues. Using high-resolution 3D imaging, simulations, cell shape and cytoskeleton manipulations, we show that planar divisions resulted from a length-limitation in mitotic-phase astral microtubules which precludes them from interacting with basal polarity, and oriented spindles from the local geometry of apical domains. Accordingly, lengthening microtubules affected spindle planarity, cell positioning and crypt arrangement. We conclude that microtubule length regulation may serve as a key mechanism for spindles to sense local cell shapes and tissue forces to preserve mammalian epithelial architecture.

## INTRODUCTION

Proliferative monolayered epithelia support the morphogenesis, function and renewal of many stem cell niches and organs during development and adult life (Godard and Heisenberg, 2019; Macara et al., 2014). They are characterized by an apico-basal polarity (A-B), and an alignment of mitotic spindles and consequent cell divisions within the tissue planes. These planar divisions position divided daughter cells side by side in the epithelial layer (Aigouy et al., 2010; Morin et al., 2007; Peyre et al., 2011) to promote tissue monolayered architecture, elongation and homeostasis (Campinho et al., 2013; da Silva and Vincent, 2007; Mao et al., 2013). Accordingly, a mis-regulation of spindle planarity has been shown to impair cell positioning and tissue integrity, and proposed to drive the emergence of dysplasia, hyperplasia, or cancer stem cell populations in stem cell niches (Bergstralh and St Johnston, 2014; di Pietro et al., 2016; McCaffrey and Macara, 2011; Morin and Bellaiche, 2011; Quyn et al., 2010; Xiong et al., 2014). To date, however, spindle orientation in epithelia has been best studied in model 2D invertebrate and vertebrate tissues, which are amenable to high-resolution live imaging and genetic manipulations of spindle associated elements (di Pietro *et al*., 2016; Morin and Bellaiche, 2011). In contrast, studying cell division orientation in complex 3D mammalian tissues and organs, relevant to human physiology and diseases has been in general hampered by the difficulty of performing advanced imaging in live animal models, and by plausible pleiotropic effects of loss of function assays (Dona et al., 2022). Therefore, addressing detailed mechanisms regulating spindle orientation in proliferating mammalian epithelial tissues remains an outstanding open endeavor.

In most animal cells and tissues, mitotic spindles are positioned and oriented from forces and torques generated by their astral microtubules (MTs) pulled by dynein motors. Dynein may be evenly distributed over the cytoplasm or cortex, a situation thought to yield length-dependent MT forces, that function to center spindles and align them with the long geometrical axis of the cell (Haupt and Minc, 2018; McNally, 2013; Wyatt et al., 2015). Dynein can also be activated at specific sub-cellular cortical polarity domains enriched in dynein activators including NuMA (<u>Nu</u>clear and <u>M</u>itotic <u>A</u>pparatus) and LGN (<u>L</u>eucine-<u>G</u>lycine-Asparagi<u>n</u>e), such as during asymmetric divisions (David et al., 2005; Morin and Bellaiche, 2011; Schaefer et al., 2000; Yu et al., 2000). In tissues, these geometrical and polarity cues often compete to dictate spindle and division positioning (Chanet et al., 2017; Godard et al., 2021; Niwayama et al., 2019; Pierre et al., 2016). For instance, the planar orientation of mitotic spindles in columnar monolayered epithelia is thought to rely on the localization of dynein-regulating polarity complexes at the level of lateral junctions, as well as on a complete mitotic rounding that serves to erase the influence of cell geometry and ensures that short mitotic-phase astral MTs bounded by dynamic instabilities reach the cortex to influence spindle orientation (Chanet *et al*., 2017; di Pietro *et al*., 2016; Lancaster and Knoblich, 2012; Luxenburg et al., 2011; Morin and Bellaiche, 2011). In general, however, how cell shape changes, MT dynamics and polarity effectors intersect to specify spindle planarity in mammalian epithelia remains poorly defined.

Given its high renewal rate throughout adult life, the intestinal tissue provides a prime model to study cell proliferation and division in mammals. The surface of the intestine is lined by a monolayer of tall columnar epithelial cells, with mitotic stem and transit-amplifying cells exclusively located in curved tissue invaginations called crypts (Gehart and Clevers, 2019; Tan and Barker, 2014). Previous studies of cell division in the crypt have reported that mitosis is associated with an apical migration of cells and DNA, which ends up with the assembly of a planar spindle in metaphase that specifies cytokinesis orthogonal to the tissue layer (Bellis et al., 2012; Carroll et al., 2017; Fleming et al., 2007; McKinley et al., 2018; Spear and Erickson, 2012). However, in part because of the lack of accessibility of the intestine *in vivo*, detailed mechanisms that control planar spindle orientation and the monolayered architecture of the crypt are still lacking.

Here, we build on intestinal organoids to address mechanisms that control division orientation in a 3D mammalian epithelium. Organoids are classically generated from isolated crypts derived from mouse intestinal pieces, and develop regular crypt-like structures with self-renewing capacity within 4 days of culture in 3D hydrogels, thereby mimicking the organization and dynamics of the *in vivo* proliferative compartment (Artegiani and Clevers, 2018; Fatehullah et al., 2016). Using live 3D imaging, physical, chemical and genetic manipulations, we show that dynein-regulating polarity complexes accumulate at the basal face of mitotic cells and not at lateral junctions, and that mitotic cells remain largely elongated along the A-B axis, challenging previous generic models for spindle planarity established in model tissues. We propose a new quantitative model for planar spindle orientation, based on a local apical cell shape sensing, mediated by length-limitation of mitotic-phase (M-phase) astral MTs. This model predicts dose-dependent variations in spindle planarity in normal organoids and in multiple conditions that affect cell shape and polarity.

## RESULTS

### Mitotic spindles in intestinal crypts orient orthogonal to the cell long axis

The spherical geometry of intestinal organoid crypts grown in 3D, provides a unique opportunity to document cell division with an optimal optical resolution along the A-B axis of the monolayer. We imaged spindle assembly and positioning along this axis, using α-tubulin-GFP/H2B-mCherry organoids (Figure 1a). We confirmed that as in many columnar epithelia, centrosomes were initially located close to the apical pole during interphase, and migrated towards the DNA at the onset of mitosis to assemble a metaphase spindle that oriented orthogonal to the A-B axis, and thus in the tissue plane (Figure 1b; Figure S1a; Videos S1 and S3) (Bellis *et al*., 2012; Carroll *et al*., 2017; McKinley *et al*., 2018). This planar orientation was maintained through metaphase, anaphase and telophase, yielding cytokinesis that bisected mother cells along the A-B axis, and placed daughter cells side by side in the monolayer (Figure 1b; Video S1) (Carroll *et al*., 2017; McKinley *et al*., 2018).

**Figure 1:**
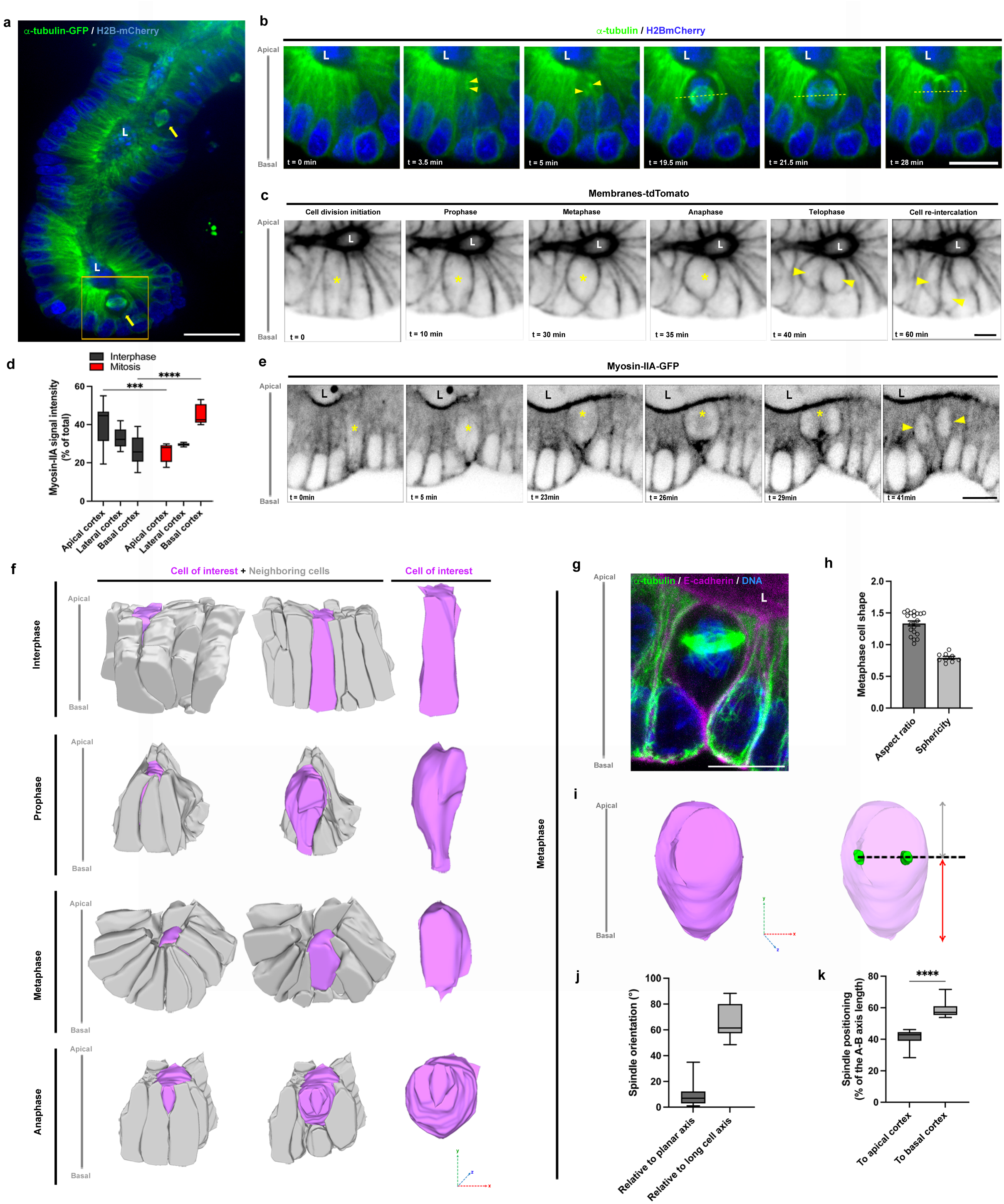
An atypical metaphase cell rounding and spindle orientation in intestinal organoids. **a** Spinning-disk analysis of α-tubulin-GFP (green) and H2B-mCherry (blue) distribution in intestinal crypt organoids. Yellow arrows point on organoid crypt-like structures. L, lumen. Scale bar, 50μm. **b** Time lapse images of α-tubulin-GFP (green) and H2B-mCherry (blue) during mitosis. Yellow arrowheads point on migrating centrosomes. Yellow dotted lines highlight spindle axis. Scale bar, 10μm. **c** Time lapse images of tdTomato organoids showing mitosis progression. Yellow star points on a dividing cell. Yellow arrowheads point on daughter cell separation and re-integration in the epithelial monolayer. Scale bar, 5μm. **d** Statistical analysis of the percentage of total signal intensity for myosin-IIA-KI-GFP at the apical, lateral or basal cortex in interphase (grey) or during mitosis (red). n = 10 cells. Two-way ANOVA test and Tukey’s multiple comparison test, ***p <0.001, ****p <0.0001. **e** Time-lapse of a live-imaged cell division in myosin-IIA-KI-GFP organoid crypt. Yellow star points on a dividing cell. Yellow arrowheads point on daughter cell separation and re-integration in the epithelial monolayer. Scale bar, 10μm. **f** Representative 3D rendering of metaphase cell (magenta) and neighboring cells (grey) after segmentation of cell membranes based on confocal z-stacks in crypt organoids. Spatial coordinates are shown. **g** Confocal analysis of α-tubulin (green) and E-cadherin (magenta) distribution in organoid metaphase cells. Scale bar, 10μm. **h** Quantitative analyses of the aspect ratio (distance between the spindle axis and the basal membrane over the distance between the spindle axis and the apical membrane) (n = 20 cells) and sphericity (n = 9 cells) of the organoid metaphase cells. Aspect ratio = 1.33±0.04 (mean±S.E.M), sphericity = 0.79±0.02. **i** Representative 3D rendering after segmentation of cell membranes and spindle poles from confocal z-stacks of organoid metaphase cells. Cell shape is depicted in magenta and in transparency, spindle poles in green. Distances between the spindle axis (black dotted line) and the apical (grey arrow) or basal membrane (red arrow) are shown. Spatial coordinates are shown. **j** Statistical analysis of the spindle orientation relative to planar axis (grey) or long cell axis (light grey). Spindle orientation relative to planar axis = 8.61±1.52° (mean±S.E.M), to long cell axis = 66.13±3.8°. N = 3 experiments, n(planar axis) = 25 cells, n(long cell axis) = 12 cells. **k** Statistical analysis of the percentage of the distance between the spindle axis and the apical (grey) or basal (light grey) membrane. Distance from apical membrane = 40.728±1.485 % (mean±S.E.M), from basal membrane = 59.272±1.485 %. N = 3 experiments, n = 13 cells. Paired t-test, ****p <0.0001.

To decipher mechanisms which regulate spindle planarity, we first explored the role of cell geometry, which generally orients spindles along the long cell shape axis (Lechler and Mapelli, 2021; Salle and Minc, 2021). We imaged cell contours in live tissues using td-Tomato organoids. This revealed that interphase columnar cells deform at the onset of prophase from the basal pole, yielding a water drop-like metaphase cell shape, rounder at the apical pole and elongated towards the basal pole (Figure 1c; Figure S1b). These mitotic shape changes were concomitant with a marked basal enrichment of myosin–IIA-GFP as well as P-MLC2, suggesting they are driven by an anisotropic actomyosin-generated cortical tension (Figure 1d-e; Figure S1c-f, Video S2). This singular actomyosin distribution may underpin mitotic apical migration and neighboring cell rearrangements on the basal side of the epithelium. In fact, the basal actomyosin clustering mirrored neighboring cell remodeling and multicellular (3-, 4- and 5-cell) contacts formation in metaphase (Figure S1g). Thus, in contrast to many epithelial tissue and adherent cells in which the actomyosin cortex homogenously remodels to ensure a complete mitotic rounding (Chanet *et al*., 2017; Luxenburg *et al*., 2011; Nakajima et al., 2013; Thery et al., 2005), intestinal organoid crypt cells exhibit an asymmetric and partial rounding and remain elongated along the A-B axis.

Full 3D reconstitution of confocal tissue z-stacks allowed to quantify metaphase cell shapes along with spindle orientations and positions (Figure 1f; Figure S1b). This showed that metaphase cells exhibit a mean aspect ratio of 1.33±0.04 along the A-B axis, and a sphericity of 0.79±0.02 (Figure 1g-i), with a mean angle between mitotic spindles and the cell’s long axis of 66.1±3.8° (Figure 1j). In addition, spindles were not centered at the geometrical cell center. Rather, they were shifted asymmetrically at an average position of 40.7±1.5 % of the cell long axis length towards the apical pole (Figure 1g,i,k). Importantly, similar partial mitotic rounding, planar and asymmetrically positioned spindles were observed *in vivo* in crypts of mouse jejunum (Figure S2a-d). Therefore, planar cell division in intestinal crypts may not directly follow long-axis geometrical rules, with a spindle oriented nearly orthogonal to the long cell shape axis and positioned asymmetrically towards the apical cell domain.

### Dynein-regulating polarity complexes localize to basal poles of mitotic cells and may not contribute to spindle orientation

These observations prompted us to assay the localization of dynein-associated polarity complexes which are potential candidates to override geometric guiding cues (Kotak et al., 2012; Morin *et al*., 2007; Niwayama *et al*., 2019; Peyre *et al*., 2011; Pierre *et al*., 2016; Zheng et al., 2010). We imaged multiple components of evolutionary conserved polarity complexes, which control spindle orientations in many tissues, including the dynein-regulators NuMA, LGN as well as LGN-binding partner afadin (Carminati et al., 2016; Wee et al., 2011). During interphase, NuMA was mostly localized within the nucleus, and re-located to spindle poles and to the cortex throughout mitosis, as reported in many systems (Compton and Cleveland, 1993; Van Ness and Pettijohn, 1983) (Figure S2e). However, although we expected the cortical pool of NuMA to localize to lateral poles in face of spindle poles, as described in most epithelia and adherent cells (di Pietro *et al*., 2016), it was largely enriched to basal poles of metaphase cells, away from and orthogonal to spindles, in both intestinal organoids and *in vivo* jejunum (Figure 2a-b; Figure S2e-f). Similar basal cortical enrichments were found for LGN and afadin (Figure 2c-d; Figure S2g-h). Dynein was in contrast localized throughout the cytoplasm and cortex, and accumulated at spindle poles (Figure S2i-j). These data show that canonical polarity complexes are segregated in an unconventional manner, away from lateral cortices and spindle planes in intestinal crypts. Rather, they appear to follow sites with increased cortical and junctional tension, as evidenced by the basal accumulation of actomyosin cytoskeleton (Figure S1c-f) and E-cadherin (Figure S2k-n) as previously reported in mammalian cell lines (Carminati *et al*., 2016; Gloerich et al., 2017; Hart et al., 2017).

**Figure 2:**
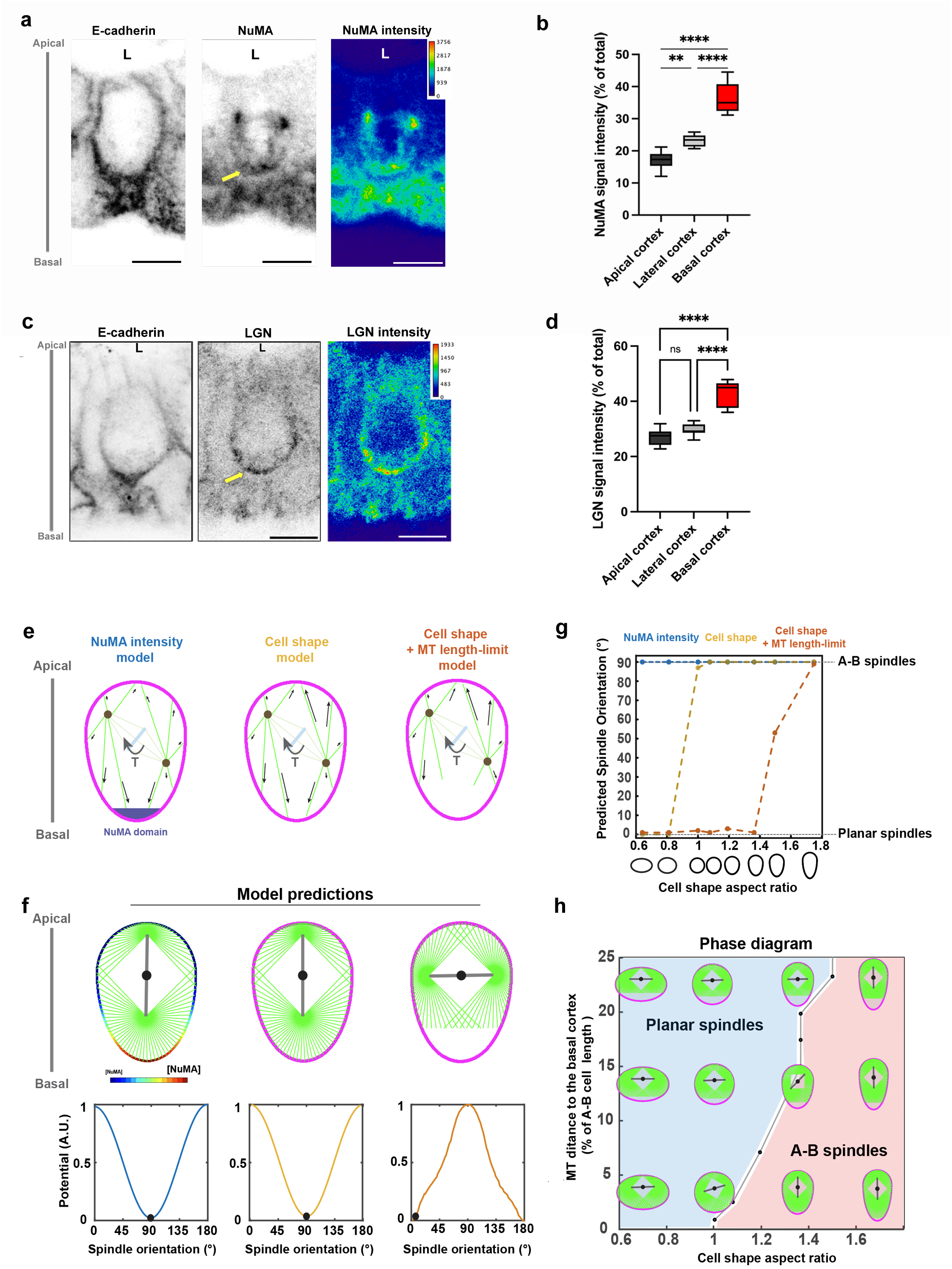
A 2D model for spindle orientation predicts that length limitation in M-phase astral microtubules may account for spindle planarity. **a** Confocal analysis of the distribution of E-cadherin and NuMA in mouse organoid metaphase cells. NuMA signal intensity is color-coded with Fire LUT table from ImageJ on the right panel. Color scale bar indicates the gray value intensity. Yellow arrow points to basal accumulation of NuMA. Scale bar, 5 μm. **b** Statistical analysis of NuMA signal intensity at the apical, lateral or basal cortex. Signal intensity at the apical cortex = 17.04±0.86 (mean±S.E.M), lateral cortex = 23.23±0.58, basal cortex = 36.26±1.5. n = 10 cells. One-way ANOVA test and Tukey’s multiple comparison test, **p = 0.008, ****p <0.0001. For each experiment, three independent experiments were carried out. **c** Confocal analysis of the distribution of E-cadherin and LGN in human organoid metaphase cells. LGN signal intensity is color-coded with Fire LUT table from ImageJ on the right panel. Color scale bar indicates the gray value intensity. Yellow arrow points to basal accumulation of LGN. Scale bar, 5 μm. **d** Statistical analysis of LGN signal intensity at the apical, lateral or basal cortex. Signal intensity at the apical cortex = 26.93±0.89 (mean±S.E.M), lateral cortex = 29.69±0.67, basal cortex = 43.14±1.42. n = 10 cells. One-way ANOVA test and Tukey’s multiple comparison test, ****p <0.0001. For each assay, three independent experiments were carried out. **e** Schemes representing the core hypothesis of the distribution of microtubule (MT) length and polarity domains in the 3 tested models for spindle orientation. **f** Rotational potential energy profiles plotted as a function of spindle orientation angles with respect to the planar axis predicted by the 3 different models. The black dot marks the preferred spindle orientation at the minimum of the potential profiles. **g** Predicted preferred spindle orientation for a range of cell shapes with increasing aspect ratios in the tissue plane or along the A-B axis. Note how the model based on cell shape coupled to length limitation in MTs transits from predicting planar spindles to apico-basal oriented spindles when cells become over-elongated, along the A-B axis. **h** Phase diagram of the predicted preferred spindle orientations for the model based on cell shape and a limit in MT lengths, drawn as a function of the 2 control parameter, the distance of MT-(+)tips to the basal side and the elongation of mitotic cell shapes.

To identify potential mechanical designs in astral microtubule (MT) force distributions that orient planar spindles in this tissue, we developed 2D mathematical models. Starting from a series of idealized cell shapes elongated along or orthogonal to the tissue plane, we placed spindles asymmetrically towards the apex, as in experiments, and computed the torque exert generated by astral MTs as a function of spindle orientation, to identify rotational equilibrium angles (Bosveld et al., 2016; Minc et al., 2011; Thery et al., 2007). As expected, when MTs were grown to fill the whole cell and exert length-dependent forces (shape-sensing system (Minc *et al*., 2011)), spindles oriented along the long cell shape axis, in the A-B axis of the tissue. Similarly, assuming that astral MT forces were scaled to the amount of the dynein-regulator NuMA (Bosveld *et al*., 2016), also oriented spindles to face NuMA basal domains along the A-B axis (Figure 2e-g). We conclude that previously established generic models for spindle orientation may not simply account for the observed spindle planarity, and that canonical polarity complexes may not influence spindle orientation in this tissue.

### Length-limitations of M-phase astral MTs mediated by the depolymerizing kinesin Kif18B promote spindle planarity

Because M-phase astral MT growth and lengths are bounded by dynamic instabilities, we tested a model based on a length limitation of astral MTs (Stout et al., 2011; Verde et al., 1992). We reasoned that such limitation could in principle prevent astral MTs from reaching basal poles of elongated mitotic cells, and create an anisotropy of mitotic aster pair shapes now longer along the planar axis. Remarkably, in 2D models, this limitation coupled to length-dependent MT forces robustly predicted planar spindle orientation for a range of cell shapes and model parameters (Figure 2e-h; Figure S3a-d). Interestingly, however as cells became too elongated along the A-B axis, spindles eventually turned to a preferred orientation along the long A-B axis, yielding a bi-stable phase diagram, controlled by the distance from MT (+)-tips to the basal poles and cell shape aspect ratio (Figure 2f-h). These modelling results suggest that a cell shape sensing mechanism truncated by a limit in MT length, could in principle function to orient spindles in the plane of intestinal organoids.

To test this hypothesis, we imaged the distribution of astral MTs around spindles. We implemented expansion microscopy of intestinal organoids to visualize and quantify individual astral MTs. This revealed the presence of astral MT (+)-tips in close contacts with both apical and lateral cortices (Figure 3a,c; Figure S3e-g). In sharp contrast, MTs were barely detected in contact with the basal side, with MT (+)-tips located at a distance ~4x higher on average from the basal cortex than the apical one (Figure 3a-c; Figure S3e,h). Importantly, these effects were not caused by a putative anisotropy in MT nucleation, as the number of MTs growing apically vs basally was similar (Figure 3d). In addition, imaging of the (+)-tip associated protein EB3-GFP confirmed this exclusion in live tissues, and indicated that the asymmetric basal exclusion of MTs was inherited from the initial apical migration of centrosomes and asters (Figure S3i, Video S3). To further test this differential MT-cortex interaction, we performed laser severing of groups of MTs in α-tubulin-GFP organoids along a line-scan orthogonal to the A-B axis, placed either on the apical or basal side of spindles (Figure 3e-f; Figure S3j-k). Both caused spindle recoils away from the cuts suggesting that astral MTs predominantly exert pulling forces, presumably by engaging with dynein motors at the cortex (Farhadifar et al., 2020; Grill and Hyman, 2005). However, in agreement with the lesser extent of MTs reaching the basal cortex, basal cuts caused significantly smaller recoils than apical ones (Figure 3f; Figure S3k). These data further support an asymmetry in MT cortex contacts along the A-B axis, and a lack of contribution of basal polarity cues to MT forces and spindle orientation.

**Figure 3:**
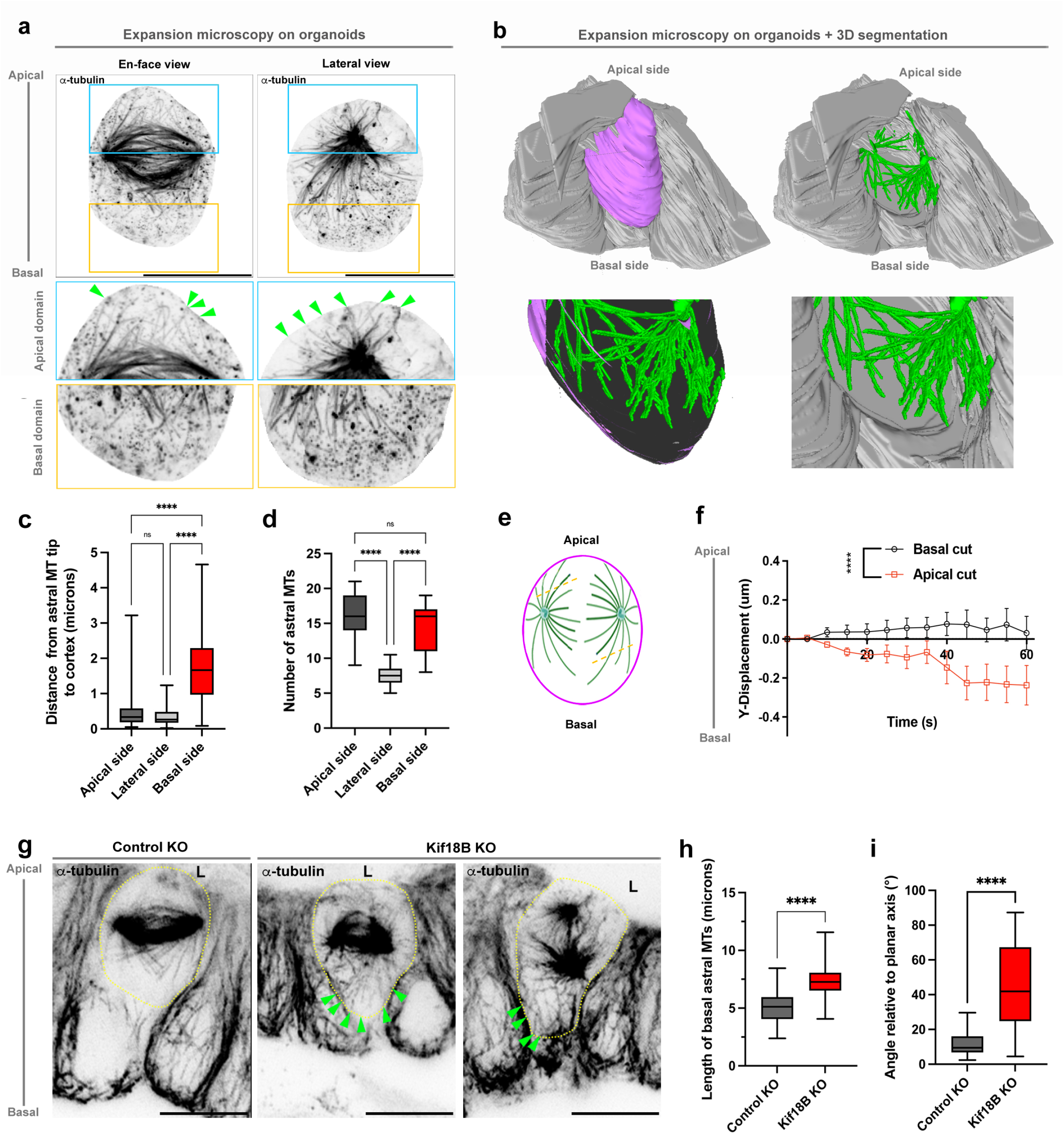
Existence of a length-limit for M-phase astral MTs that impact spindle orientation. **a** Projected confocal z-stacks of α-tubulin distribution in isolated organoid metaphase cell. En-face and lateral views are presented. Apical domain areas boxed in blue and basal domain areas boxed in yellow are presented in the low panels. Green arrowheads point to astral MTs contacting the apical cortex. Corrected scale bar, 10μm. **b** Representative 3D rendering of a metaphase cell (magenta), neighboring cells (grey), spindle poles and microtubules (green) after segmentation of a confocal z-stack of E-cadherin and α-tubulin localization. Are depicted A-B views tilted to show basal cortex. **c** Statistical analysis of the distance between the astral MT tip and the apical, lateral or basal cortex. Distance between astral MT tip and the apical cortex = 0.466±0.045 (mean±S.E.M), the lateral cortex = 0.349±0.025, the basal cortex = 1.685±0.086. N = 3 experiments, n = 11 cells. N = 104 apical astral MTS, n = 102 lateral astral MTs, n = 107 basal astral MTS. Two-way ANOVA with Tukey’s multiple comparisons test, ****p <0.0001; ns, non-significant. **d** Statistical analysis of the number of astral MTs at the apical, lateral and basal sides. Number of astral MTs at the apical side = 16.3±1.09 (mean±S.E.M), at the lateral side = 7.6±0.5, at the basal side = 14.2±1.06. N = 3 experiments, n = 11 cells. Unpaired t-test, ****p <0.0001; ns, non-significant. **e** Scheme showing the position of apical or basal laser cut (yellow dotted lines) of astral MTs. **f** Statistical analyses of the spindle pole displacement in the y-axis after apical or basal laser cut in organoid metaphase cells. n = 6 cells for each condition. Unpaired t-test, ****p <0.0001. **g** Confocal analysis of α-tubulin distribution in metaphase cells in control-KO or Kif18B-KO organoids. Yellow dotted line delimits the metaphase cell. Green arrowheads point to astral MTs contacting the basal cortex. L, lumen. Scale bar, 10μm. **h** Statistical analysis of the length of basal astral MTs in metaphase cells of control-KO or Kif18-KO organoids. n = 99 astral MTs in 13 control-KO cells, n = 103 astral MTs in 17 Kif18-KO cells. Unpaired t-test, ****p <0.0001. **i** Statistical analysis of the spindle axis orientation relative to the planar axis of the epithelium in metaphase cells of control-KO or Kif18-KO organoids. n = 32 control-KO cells, n = 50 Kif18-KO cells. Unpaired t-test, ****p <0.0001. For each experiment, three independent experiments were carried out.

To directly assay the role of MT length regulation, we next sought to increase astral MTs length and assess impact on spindle orientation. Kif18B, a (+)-tip depolymerizing kinesin, was shown to promote MT catastrophe and thus limit the length of M-phase astral MTs in mammalian cells and tissues (McHugh et al., 2018; Stout *et al*., 2011). Accordingly, knocking out Kif18B, using a CRISPR-Cas9 inducible system in organoids, led to significantly longer astral MTs, so that even the basal-facing MTs now grew long enough to reach the cell cortex (Figure 3g-h; Figure S3l-m). Strikingly, spindles in Kif18B-KO organoids were not planar anymore, and rather oriented randomly, with a significant fraction of spindles oriented along the A-B axis (Figure 3i). Together, these results suggest that a length-limitation of mitotic astral MTs mediated in part by depolymerizing kinesins, may prevent MTs from interacting with basal polarity factors and allow spindles to orient in the tissue plane by probing the geometry of the apical fraction of dividing cells.

### A 3D model for spindle position and orientation in intestinal crypts

In order to validate this length-limitation hypothesis, we explored the parameter space in the model phase diagram (Figure 2f,h) by experimentally altering cell shapes (Figure 4a-c). We first elongated cells in the tissue plane, to validate a general influence of apical cell shape. We grew organoids in L-WNR medium, an exogenous global Wnt3a treatment, which results in the formation of hyper-proliferative and undifferentiated cystic structures (Farin et al., 2012; Sato et al., 2011b), in which the epithelial monolayer displays a flat squamous morphology with cells now elongated in the tissue plane (Figure S4a). As predicted by models, and not surprisingly, spindles oriented parallel to the long cell shape axis in the tissue plane in this condition (Figure 4a-d; Figure S4a). To increase mitotic cell elongation along the A-B axis, we next affected contractility by inhibiting myosin-IIA activity. Both blebbistatin treatment or myosin-IIA-KO drastically impaired mitotic shape changes and rounding, with cells remaining significantly more elongated along the A-B axis than controls (Figure 4a-c; Figure S4b-e). This resulted from a loss of cellular rearrangements and the formation of fewer multicellular contacts in the basal environment of metaphase cells (Figure S4d-h). These results confirmed the primary function of basal actomyosin contractility in reshaping mitotic cells and its contacts with neighbors in this tissue. Importantly, as predicted in models (Figure 2f,h), in these more A-B elongated metaphase cells, spindles now turned to preferentially orient along the long A-B cell shape axis (Figure 4a-b,d). Importantly, in these conditions, polarity complexes detached from the cell cortex (Figure S4i-j) and spindles were still positioned off-center towards the apical cell poles (Figure 4b), ruling out putative contributions of basal polarity to this A-B spindle orientation.

**Figure 4:**
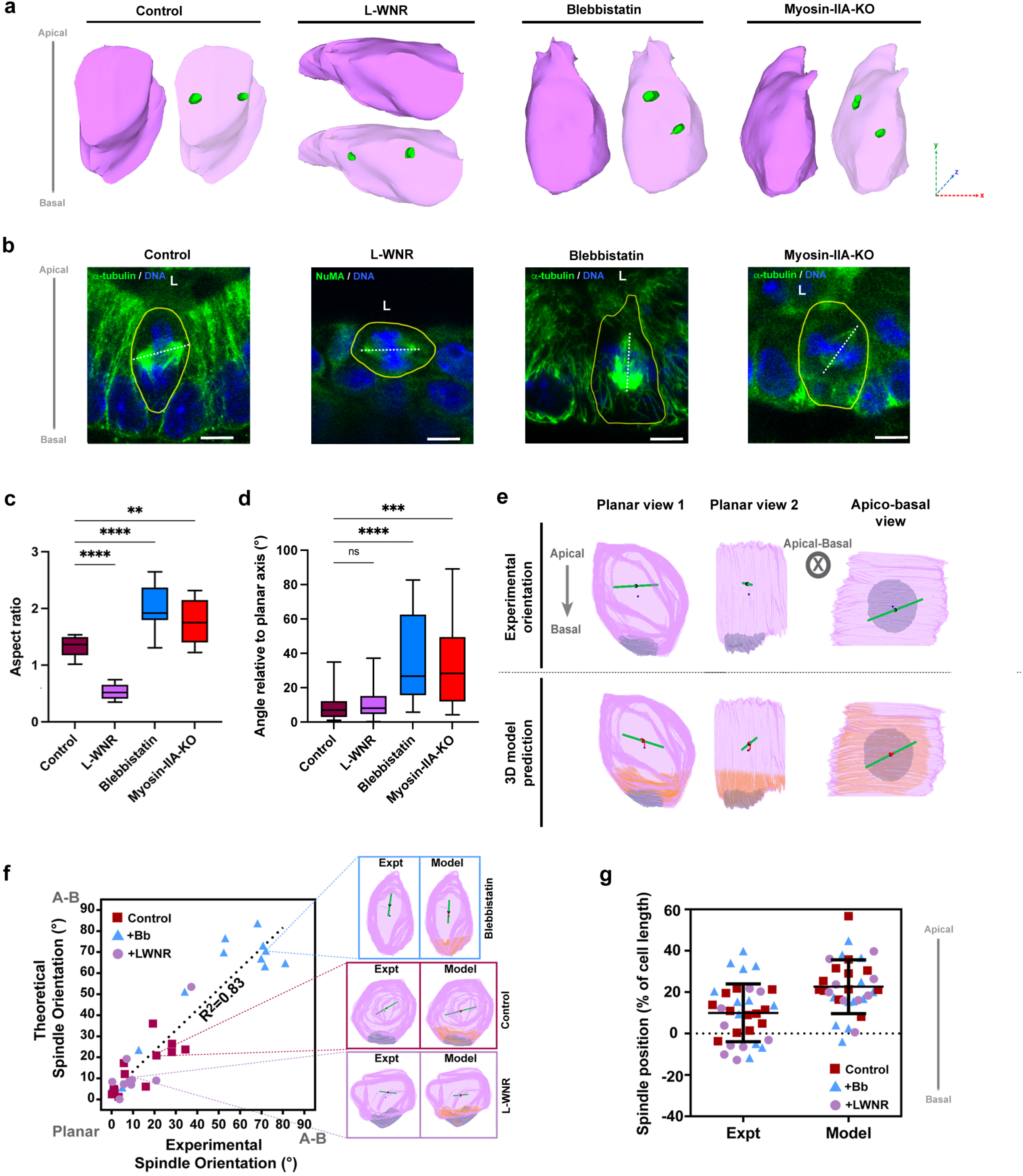
Metaphase cell shape impacts planar spindle orientation in organoid cells. **a** Representative 3D rendering of cell shape and spindle pole positioning after segmentation of cell membranes and NuMA spindle-associated signal from confocal z-stacks of control (DMSO-treated), L-WNR-cultured, blebbistatin-treated or induced myosin-IIA-KO organoid metaphase cells. Cell shape is depicted in magenta, spindle poles in green. Spatial coordinates are shown. **b** In control (DMSO-treated), L-WNR-cultured, blebbistatin-treated or induced myosin-IIA-KO organoid metaphase cells, is shown the confocal image of α-tubulin or NuMA (green) and nuclei (blue) Scale bar, 5μm. **c** Statistical analyses of organoid metaphase cell aspect ratio in control (DMSO-treated), L-WNR-cultured, blebbistatin-treated or induced myosin-IIA-KO organoid metaphase cells. Aspect ratio of metaphase cells in control organoids = 1.33±0.04 (mean±S.E.M), in L-WNR-cultured organoids = 0.53±0.04, in blebbistatin-treated organoids = 2.01±0.12, in myosin-IIA-KO organoids = 1.78±0.13. n (control organoids) = 20 cells, n (L-WNR organoids) = 10 cells, n (blebbistatin-treated organoids) = 10 cells, n (myosin-IIA-KO organoids) = 10 cells. One-way ANOVA, with Tukey’s multiple comparisons tests, **p = 0.0011, ****p <0.0001. **d** Statistical analyses of the spindle axis orientation relative to the plane of the epithelium. Angle deviation in control organoids = 8.61±1.52 (mean±S.E.M), in L-WNR-cultured organoids = 11.28±3.74, in blebbistatin-treated organoids = 37.11±5.04, in myosin-IIA-KO organoids = 34.88±5.22. n (control organoids) = 25 cells, n (L-WNR organoids) = 9 cells, n (blebbistatin-treated organoids) = 25 cells, n (myosin-IIA-KO organoids) = 25 cells. One way-ANOVA, ***p = 0.0001, ****p <0.0001. ns, non-significant. For each experiment, three independent experiments were carried out. **e** (Top) 3D reconstitution of experimental cells obtained from 3D z-stacks of a representative dividing cell labeled for E-cadherin and NuMA to extract cell shape (magenta), spindle orientation (green line) and NuMA accumulation at the basal cortex (grey domain). (Bottom) Corresponding simulation output, based on MT length-dependent forces (shape sensing) and an exclusion of MTs length in the basal domain region (orange zone). The red traces represent the history of spindle axis displacement in the simulation. Both experimental and simulated cells are viewed in different planes with respect to A-B axis. **f**. Predicted spindle orientation angle with respect to the A-B axis, plotted as a function of the experimental axis for 10 individual control, L-WNR treated or blebbistatin-treated crypt cells. Insets provide representative examples of experimental and simulated spindle orientation in the different treatments. The dotted line is a linear fit, with a slope of 0.96 (1 being a perfect agreement between models and experiments). R^2^ is the correlation coefficient between the fit and the data. **g**. Experimental and theoretical prediction of spindle asymmetric position towards the apical cell poles in the indicated conditions.

To test our hypothesis in these multiple conditions affecting cell shapes using realistic 3D cell geometries, we next turned to simulations. We adapted a gradient descent strategy that can predict both spindle orientation and position in 3D from the 3D geometry of mitotic cells, the localization of polarity cues and length limitation in astral MTs (Pierre *et al*., 2016). Using 3D segmentations of individual cell contours, spindle poles and basal domains from fixed tissues, we reconstituted experimental cell shapes, polarity and spindle orientation and position in mitotic cells (Figure 4e). By inputting the experimental geometry and a limit for MT growth at the basal pole in the model, we ran simulations starting from random positions and orientations, and searched for equilibrium. The simulations were robust to initial conditions and predicted with accuracy in 3D, the planarity of spindles and their apical shifts in control and L-WNR, as well as their reorientation along the A-B axis in more elongated blebbistatin-treated cells (Figure 4e-g; Figure S5a-c, and Videos S4-5). In contrast, models in which astral MTs filled whole cell volumes, or models based on a predominant influence of NuMA domains, yielded poor agreements with experiments (Figure S5d-g). Importantly, the model predicted deviations to planarity in a dose-dependent manner, among more or less elongated wt or blebbistatin treated cells, and also the correct orientation of spindles within the tissue plane, when shape anisotropies were present in this plane (Figure 4e-f). These direct comparisons between 3D model predictions and experiments strongly support that length-limitations of astral MTs can account for spindle position and planar orientation in intestinal crypts.

### Lengthening astral MTs affect epithelial tissue layering

This truncated shape sensing may have many advantages for the regulation of tissue layering and cell density. For instance, it may allow cells to adapt division orientation to planar tissue forces, tilting spindles along the A-B axis if cells become over-compressed by their neighbors, providing a potential homeostatic mechanism to regulate cell density and monolayered architecture (Wyatt *et al*., 2015). Accordingly, in Kif18B-KO organoids, as a consequence of spindle mis-orientation, planar polarity of cytokinesis and placement of daughter cells were largely impaired (Figure 5a-b). Hence, the regular basal nuclear arrangement, which is a hallmark of polarized columnar epithelia, was lost three days after KO induction (Figure 5a,c-d). Finally, nuclei densities were significantly reduced in the Kif18B-KO (Figure 5e), showing how spindle mis-orientation may impact crypt architecture and cell density. These results directly demonstrate how a modulation in astral MT length may impact spindle planarity and architecture of a mammalian tissue.

**Figure 5:**
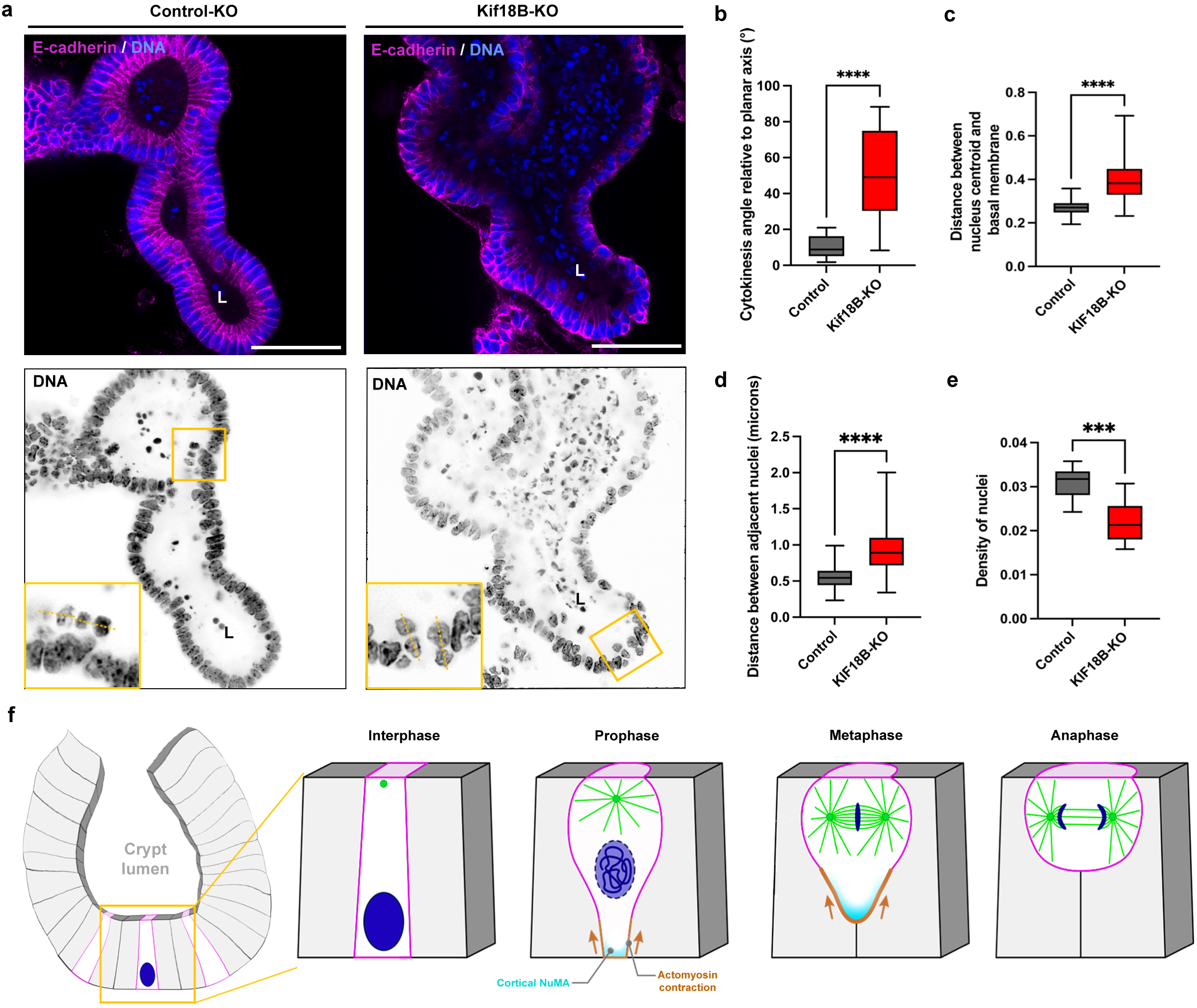
Impact of spindle mis-orientation in epithelial organization. **a** Confocal microscopy analysis of the distribution of the E-cadherin (magenta) and nuclei (blue) in control-KO and Kif18B-KO organoids. Representative cytokinesis events are boxed in yellow and presented in bottom left. Yellow dotted lines highlight cell division axis. L, lumen. Scale bar, 50μm. **b** Statistical analysis of cytokinesis angle relative to the planar axis in control-KO or Kif18B-KO organoids. n = 13 control-KO cells, n = 26 Kif18B-KO cells. Cytokinesis angle (control-KO) = 10.3±1.68°, cytokinesis angle (Kif18B-KO) = 50.8±4.9°. Unpaired t-test, ****p < 0.0001. **c** Statistical analysis of interphase nuclear positioning (distance from the centroid of the interphase nucleus to the basal membrane) (n = 100 cells). Nuclear positioning (control-KO) = 0.271±0.003 μm (mean±S.E.M), (Kif18B-KO) = 0.399±0.01 μm. Unpaired t-test, ****p < 0.0001. **d** Statistical analysis of distance between adjacent interphase nuclei (n = 100 cells). Distance between nuclei (control-KO) = 0.559±0.015 μm (mean±S.E.M), (Kif18B-KO) = 0.939±0.032 μm. Unpaired t-test, ****p < 0.0001. **e** Statistical analysis of nucleus density (n = 10 fields). Nucleus density (control-KO) = 0.031±0.001 μm (mean±S.E.M), (Kif18B-KO) = 0.022±0.001 μm. Unpaired t-test, ***p = 0.0002. **f** Scheme depicting the proposed model of spindle polarity modulation in organoid metaphase cells.

## DISCUSSION

### A new generic model for planar spindle orientation in proliferative epithelia

How spindle orientation is regulated in proliferative tissues and stem cell niches is of fundamental importance for organ morphogenesis and homeostasis, and highly relevant to human disorders (Caussinus and Gonzalez, 2005; McCaffrey and Macara, 2011; Nakajima *et al*., 2013; Seldin and Macara, 2017). Here, by documenting with high temporal and spatial resolution spindle orientation together with MT dynamics, cell shape and polarity in a 3D mammalian proliferative organoid, we propose a new model for the control of planar spindle orientation and monolayered tissue architecture. This model is based on a partial cell shape sensing resulting from a length-limitation of M-phase astral MTs. This limitation is coupled to the A-B polarity of the columnar tissue, and allows initially apically positioned centrosomes to stop their basal migration at a position shifted towards the apex, and astral MTs to probe the local apical fraction of mitotic cells to orient spindles in the tissue plane (Figure 5f). This mechanism has similarities with previous models proposed for asymmetric aster pair positioning and orientation in some large zygotes featuring anisotropic MT asters (Pierre *et al*., 2016; Wuhr et al., 2008) (Figure 5f). Importantly, our model contrasts with established ones in many epithelia, in which the role of polarity effectors including NuMA, LGN or afadin recruited to lateral cortices, is thought to be predominant to orient spindles in the tissue plane (di Pietro *et al*., 2016). In intestinal crypts, their unconventional localization to the basal domain of mitotic cells suggest they may not contribute to orient spindles, and raise the question of how they may be segregated there. We found that myosin-IIA inhibition caused their detachment from the basal cortex (Figure S4i-j). Therefore, we suggest that basal polarity recruitment could be driven by enhanced local cortical tension associated to basal actomyosin activity and cellular rearrangements (Figure 1d-e; Figures S1a-g), as proposed in epithelial cell lines (Bosveld *et al*., 2016; Carminati *et al*., 2016; Gloerich *et al*., 2017; Hart *et al*., 2017). Other plausible mechanisms contributing to this basal localization might include a local clearance of polarity effectors from the apex from chromosome-derived signals (Dimitracopoulos et al., 2020; Kiyomitsu and Cheeseman, 2012), or a recruitment associated to the numerous basal multicellular contacts formed in mitosis (Bosveld *et al*., 2016).

Interestingly, asymmetric actomyosin activity in other epithelial cells and tissues has been proposed to directly influence spindle orientation. In drosophila, one study notably reported that local actomyosin enrichment may override geometrical cues to orient spindles in face of actomyosin rich zones (Scarpa et al., 2018). In cultured cells, a recent local optogenetic assay showed an opposite behavior with spindles orienting orthogonal to the most contractile cortical zone (Kelkar et al., 2020). Our findings in the crypt align more with this latter finding, at least in term of geometrical outputs. However, our results favor a more indirect role for basal actomyosin asymmetric enrichment, primarily impacting spindle orientation through its role in driving a partial and asymmetric mitotic rounding. We currently do not understand which mechanism may enrich actomyosin at the basal poles of mitotic crypt cells. We suggest it may reflect the need for apical migrations of dividing cells in columnar tissues, as proposed in the mouse nervous system or zebrafish retina (Norden et al., 2009; Schenk et al., 2009). Importantly, in addition to promote anisotropic rounding, we also propose that this pool functions to maintain epithelial cohesion at the base of mitotic cells (Figure 1d-e; Figures S1a-g, 4d-f). Further exploration of the regulation and role of this mitotic basal actomyosin enrichment, may reveal important aspects of intestinal crypt dynamics and architecture.

### Length-regulation of M-phase MTs, mitotic rounding and spindle orientation

Astral MTs control spindle positioning in organisms ranging from yeast to mammals (McNally, 2013). While most models have thus far largely assumed that astral MTs grow to reach the cell cortex, to probe cell shapes and polarities, recent studies have suggested that M-phase astral MTs which are bounded by dynamic instabilities, may be just long enough in mitosis to reach the cortex of fully rounded cells (Mitchison et al., 2015). Accordingly, impairment of mitotic rounding has been shown to prevent MTs from properly reaching and interacting with the cortex, yielding defects in chromosome segregation and spindle orientation (Lancaster et al., 2013; Luxenburg *et al*., 2011). In our study, we propose that such cell-cycle regulated length limitation is directly exploited by the tissue to allow the spindle to only probe apical cell shapes, position asymmetrically and orient in the tissue plane. Results from Kif18B-KO organoids favor a global regulation of astral MT catastrophe and length in mitosis, which translates into an A-B asymmetric MT cortex contact from the initial apical location of centrosomes. However, we also envisage that more local basal regulation of MT dynamics or dynein activity could emerge directly from enhanced basal actomyosin activity, or from putative higher basal organelle crowding, as proposed in other systems (Jimenez et al., 2021; Pierre *et al*., 2016; Zhu et al., 2010). While the impact of cell shape on division orientation was first reported more than 150 years ago (O., 1884), mechanisms by which geometries are being sensed in multicellular tissues are still at their infancy (Haupt and Minc, 2018). Our study provides an important new generic mechanism to orient spindles along the short axis in mammalian epithelia, and calls for a better exploration of mechanisms regulating astral MT length and mitotic shape changes in multicellular tissues.

### Functions of planar divisions for crypt dynamics and architecture

Planar cell divisions are key for maintaining the monolayered architecture of certain epithelial tissues. Conversely, programmed alterations of planar divisions along the A-B axis of the tissue are required for epithelial stratification such as during skin development in the mouse embryo (Biggs et al., 2020; Lechler and Fuchs, 2005). In the curved 3D geometry of intestinal crypts, we found that cell divisions orient parallel to the local plane of curvature of the tissue. We propose that such organization of cell divisions may contribute to promote a near-isotropic expansion of the tissue while maintaining a regular monolayered architecture. Accordingly, altering spindle planarity resulted in mis-placed daughter cells, often protruding into the apical lumen (Figure 5a-e). Whether such defects could lead to the long-term formation of tumor masses, epithelial dysplasia or hyperplasia, or if mechanisms such as apoptosis or daughter cell reintegration may safeguard the intestinal tissue, remain to be tested (Lough et al., 2019). In addition, many of the dividing cells in the crypt are Lgr5+-stem cells, which may undergo symmetric fate divisions for self-renewal or asymmetric divisions to generate a daughter cell that becomes fated. As such, whether alteration of planar spindle positioning affects the auto-renewal properties of organoids and their homeostasis, as previously described for instance in Apc-mutated intestinal tissues, is another important open question (Dow et al., 2015; Feng et al., 2013). More work on the regulation and function of oriented division in mammalian stem cell niches, tissues and organs may help to better appreciate the emergence of human epithelial disorders.

## Acknowledgments

We thank René-Marc Mège and Benoit Ladoux (Institut Jacques Monod, IJM) for helpful discussions as well as Paula Martin Gil, Valeria Rocchi and Jeremy Sallé (IJM) for technical help. We thank Renata Basto, Ana-Maria Lennon-Dumenil, Robert S. Adelstein and Danijela Vignjevic for providing mice. Confocal microscopy and spinning-disc analyses were performed in the ImagoSeine microscopy facility (Institut Jacques Monod, IJM). This work was supported by grants from the Groupama Foundation – Research Prize for Rare Diseases 2017 (to D.D), the Fondation pour la Recherche Médicale (FRM) (to C.G. and D.D.), the Human Frontier Science Program (RGP0038/2018) (to D.D), the LabEx “Who Am I?” #ANR-11-LABX-0071 and the Université de Paris IdEx #ANR-18-IDEX-0001 funded by the French Government through its “Investments for the Future” program (to N.M. and D.D.), and the INCA PLBIO20-150, Cancéropole Ile-de-France (to N.M. and D.D.), and grants from La Ligue Contre le Cancer (EL2021.LNCC/ NiM).

## Supplemental Information

Figures S1-S5 and Videos S1-S5 are available in the online version of the paper.

## Author contributions

J.S, O.F, M.A.F, M.S., C.G., T.D., HC, and D.D. performed experiments. N.M. designed and performed models and simulations. J.S, M.A.F., C.G., N.M. and D.D. designed the experiments. J.S, M.A.F, C.G., M.S., H.C., N.M. and D.D. performed analyses. J.S, N.M. and D.D. coordinated the overall research and experiments, and wrote the manuscript.

## STAR METHODS

### Resource availability

#### Lead contact

Further information and requests for resources and reagents should be directed and will be fulfilled by the lead contacts, Delphine Delacour and Nicolas Minc.

#### Materials availability

Materials generated in the current study are available from the lead contacts upon request.

#### Data and code availability

New generated codes and data obtained in the current study are available from the lead contacts on request.

### Experimental model and subject details

#### Organoid culture and transfection

Wild-type C57/Bl6 mice were provided by the animal house facility of the Institut Jacques Monod. Mice used for intestinal crypt isolation were between 6 and 12 weeks old. After euthanization by cervical dislocation, the small intestine was harvested, flushed with PBS to discard luminal content and cut longitudinally open. The tissue was then cut into small pieces of 3-5 mm and further washed in PBS. The pieces of intestinal tissue were then incubated on ice for 10 min in a tube containing 5 mM EDTA. The tube was then vortexed for 2 min to release villi from the tissue. After EDTA removal, the intestinal pieces were placed in cold PBS and vortexed vigorously for 3 min to ensure crypt release. This process was repeated 3 times, with each fraction recovered. The third and fourth fractions are usually concentrated in crypts, so these are combined and passed through a 70-μm cell strainer to remove remaining villi and centrifuged at 1000 RPM for 5 min. The pellet (crypts) was then washed in advanced DMEM/F12 (#12634010 Thermofisher Scientific, Waltham, Massachusetts, USA) and centrifuged. The final pellet is resuspended in 50 μl of 1:1 ratio of advanced DMEM/F12 and ice-cold Matrigel (#734-1100 VWR, Radnor, PA, USA) and plated as domes. Incubation at 37°C for 20-30 min allowed Matrigel polymerization. Organoid culture was performed in IntestiCult™ Organoid Growth medium (#06005 STEMCELL Technologies, Vancouver, Canada), from here on termed ENR medium. Organoids were routinely grown in Matrigel with IntestiCult™ Organoid Growth medium and passaged every 7 to 10 days. Medium was changed every 2 days. Live-imaging or immunofluorescence experiments were performed on 3-4 days organoids. For cystic growth, intestinal organoids were cultured with L-WNR medium supplemented with 10μM CHIR99021 for 10 days. The L-WNR cell line was purchased from ATCC (ATCC CRL-3276™). The L-WNR medium was produced according to the ATCC recommendations.

α-tubulin-GFP and EB3-RFP expression was carried out by lentiviral transduction using the commercial lentiviral biosensors LentiBrite (#17-10206; #17-10222, Merck, Darmstadt, Germany) and following an already established protocol for lentiviral transduction in intestinal organoids (Van Lidth de Jeude et al., 2015). This process involves dissociating intestinal organoids into single cells using TrypLE Express (#12605010 Thermofisher Scientific, Waltham, Massachusetts, USA) and seed them on a layer of Matrigel along the lentiviral vectors overnight, before covering with another layer of Matrigel the next day to allow growth into 3D organoids. The medium was enriched with 10 μM of CHIR99021 and 10 μM of Y-27632 (#72054 and #72304 STEMCELL Technologies, Vancouver, Canada) to prioritize stem cell proliferation and improve single cell survival respectively. After 48 to 72h, fluorescent organoids start appearing and can be isolated and expanded.

H2B-mCherry organoids were generated from H2B-mCherry-knock-in mice provided by Renata Basto (Institut Curie, Paris). VillinCreERT2-tdTomato organoids were generated from mice provided by Danijela Vignjevic (Insitut Curite, Paris) (Krndija et al., 2019). Myosin-IIA-GFP-knock-in mice (Zhang et al., 2012) were provided by Robert S. Adelstein (NHLBI, Bethesda) and Ana-Maria Lennon-Dumesnil (Institut Curie, Paris). Myosin-IIA-KO/mTmG mice (Krndija *et al*., 2019) were kindly provided by Danijela Vignjevic (Insitut Curite, Paris) and generated by crossing Myosin-IIA-KO (Jacobelli et al., 2010) and mTmG mice (Muzumdar et al., 2007). Cre recombinase for Myosin-IIA-KO was induced with 100nM of 4-hydroxytamoxifen for 24h (#SML1666, Sigma-Aldrich).

Kif18B-KO organoids were generated by a CRISPR-Cas9 strategy using the Edit-R All-in-one lentiviral system from Horizon Discovery (Cambridge, UK). A set of 3 lentiviral sgRNAs were used to target Kif18B (#GSGM11839-246783580: GGTCAGAACACCCAGTTAAT; #GSGM11839-246783577: GTGTTTGCCTATGGCGCCAC; #GSGM11839-246783579: GTGGTGTTGAGGTCCCGAGT). Control-KO organoids were generated using a non-targeting sgRNA (#GSG11811, Horizon Discovery). Organoids were transduced with lentiviral particles containing an inducible Cas9 along the sgRNAs for the targeted gene. Selection was then performed using 2ug/ml of puromycin (#A1113803, Thermofisher Scientific) for at least 2 weeks before inducing Cas9 with 600 ng/ml of doxycycline (#D9891 Merck).

For human organoid preparation, the project was approved by the Scientific Committee of the tumor bank of Lille and the Department of Pathology of the Lille University Hospital. The patient had signed an informed consent. Left colectomy was carried out for a colon adenocarcinoma in the Department of General and Digestive Surgery of the Lille University Hospital. The patient had not been treated with neoadjuvant chemotherapy. The normal mucosa sample (1 cm2 harvested >10cm distant from the tumor) was cut into small pieces (<3mm3) and washed thoroughly with 1X PBS buffer supplemented with combined antibiotics (normocin, gentamicin and amphotericin B). Mucosa fragments were then incubated in 25 ml of 1X PBS with 2.5mM EDTA at 4°C for 30 min under slow rotation. Mechanical release of colonic crypts was then performed three times in 10 mL 1X PBS. Fractions of colonic crypt suspension were pooled, centrifuged, resuspended in advanced DMEM/F12 medium and filtered through a 70-μm cell strainer. After counting, cells were resuspended in Matrigel and seeded in 40 μL domes in the wells of a 24-well plate. After Matrigel solidification, domes were covered with complete colon organoid medium (advanced DMEM/F12 medium supplemented with 1X Glutamax (Invitrogen), 1 X HEPES (Sigma-Aldrich), B-27® Supplement Minus Vitamin A (Invitrogen), 1 X N2 (Invitrogen), 1 mM N-acetyl-L-cysteine (Sigma-Aldrich), 50% v/v Wnt3a/RSPO1/Noggin-conditioned medium, 50 ng/ml EGF (Peprotech), 0.5 μM A83-01 (Tocris), 10 mM Nicotinamide (Sigma-Aldrich), 10 nM Gastrin (Sigma-Aldrich) and 10 μM Y-27632 dihydrochloride (Tocris), as recommended in (Sato et al., 2011a). Resulting colonic organoids were named COL-2920xi. Complete medium without Y-27632 was then renewed every two days and organoids were passaged through mechanical disruption every week.

### Method details

#### Antibodies and reagents

Mouse monoclonal antibody directed against α-tubulin (DM1A clone, IF dilution, 1:100) was purchased from Sigma-Aldrich. Rabbit polyclonal antibody directed against EpCAM (#ab71916, IF dilution, 1:100) and rabbit polyclonal antibody directed against α-tubulin (#ab18251, IF 1:100) were from Abcam. Mouse monoclonal antibody directed against E-cadherin (clone 36, #610181, IF dilution, 1:50) was from BD Biosciences. Rabbit polyclonal antibody directed against Phospho-Myosin Light Chain 2 (Ser19, #3671, IF dilution, 1:100) and rabbit monoclonal antibody directed against E-cadherin (clone 24E10, #3195S, IF dilution 1:100) were from Cell Signaling Technology. Rabbit monoclonal antibody directed against NuMA (EP3976, #ab109262, IF dilution, 1:100) was from Abcam. Rabbit polyclonal antibody directed against non-muscle myosin heavy chain II-A antibody (poly19098 clone, #909801, WB dilution 1:500) was from Biolegend. Mouse monoclonal antibody directed against GAPDH (#60004-1-Ig, WB dilution 1:5000). Phalloidin-Alexa488, 568 or 647 were from Life Technologies. Nuclei were stained with Hoechst 33342 solution incubation (Life Technologies, Paisley, UK) at a 1:1000 dilution. Blebbistatin and Y-27632 were from Sigma Aldrich (Saint-Louis, MO, USA).

#### Biochemical analysis

For western blots, organoid lysates were prepared 3 days after plating using per condition 6 wells of a 24-well plate. Matrigel was depolymerized by incubating with 1 ml of Gentle Cell Dissociation Reagent (#07174 STEMCELL Technologies, Vancouver, Canada) for 30 min at 4°C and centrifugation for 5 min at 500 x g at 4°C. The pellet was resuspended in lysis buffer containing 25mM Tris / 5 mM NaCl / 1mM EDTA / 1mM EGTA / 0.5% NP40 / 1% Triton TX100, and incubated on ice for 30 min. The solution was then passed 10 times through a syringe equipped with a 23G needle and centrifuged at 10 000 RPM at 4°C for 10 min. Supernatant total protein content was measured by Bradford assay (Biorad). For each condition, 50μg of proteins was loaded per well in Novex Tris-Glycine pre-cast gels (ThermoFischer Scientific). Proteins were detected with either HRP-linked goat anti-mouse IgG antibody (dilution 1:10,000; Sigma-Aldrich) or HRP-linked donkey anti-rabbit IgG antibody (dilution 1:10,000, GE Healthcare, Buckinghamshire, UK), and visualized on ImageQuant LAS4000 (GE-Healthcare). Signal quantification was performed using Fiji software.

#### Immunostaining

Routinely, organoids were fixed using 4% paraformaldehyde for 30 min, then permeabilized using 0.025% saponin solution in PBS for 30 min. Blocking step was performed in 0.025% saponin/1% BSA solution for 45 min, before proceeding to incubation with primary antibody at 4°C overnight. The next day, the primary antibody was removed and the organoids washed 3 times in PBS for 10 min each, before adding the secondary antibody and left to incubate for 2h at room temperature. Finally, organoids were washed 3 times again for 10 minutes before incubating in Hoechst 33342 for 15 min to stain nuclei. Immunostained samples were mounted in home-made Mowiol solution.

For the MT immunostaining, we used an established protocol that maintain the MT integrity in intestinal organoids (Goldspink et al., 2017). Briefly, organoids were isolated from the Matrigel and fixed in a methanol/formaldehyde solution (92% methanol, 8% formaldehyde). The blocking step was performed by incubating organoids in 10% goat serum solution in PBS with 0.1% Triton X-100. Immunostained organoids were mounted in home-made Mowiol solution on a slide.

For E-cadherin immunostaining *in vivo*, mouse jejunum was processed as previously described (Salomon et al., 2017). Briefly, samples were fixed for 2 hours in 4% PFA and paraffin embedded. 5 μm tissue sections were de-waxed in a xylene bath, rehydrated in isopropanol and in solutions with decreasing ethanol concentrations. Tissue sections were then blocked in 10% goat serum (Sigma-Aldrich) for 1 h. Primary antibody incubation was performed at 4°C overnight and secondary antibody incubation at room temperature for 2 h, both in 1% goat serum solution. Hoechst33342 staining was used to detect nuclei. Tissue sections were mounted in home-made Mowiol 488 solution.

For NuMA immunostaining *in vivo*, 1-mm pieces of mouse jejunum were fixed in 4% PFA overnight under shaking. After PBS wash, tissue permeabilization was performed in 1% Triton X-100 / PBS solution for 1 h, before saturation in 1% BSA / 3% goat serum / 0.2% Triton X-100 / PBS solution for 1 h. Incubation with primary or secondary antibodies were done in 0.1% BSA / 0.3 % goat serum / 0.2 % Triton X-100 / PBS overnight at 4°C. Hoechst33342 staining was used to detect nuclei. Immunostained samples were mounted in Vectashield (Vector Laboratories, Burlingame, CA).

#### Live imaging

Dynamics experiments on α-tubulin-GFP/H2B-mCherry, myosin-IIA-KI-GFP or EB3-RFP organoids were performed using an inverted Zeiss microscope equipped with a CSU-X1 spinning disk head (Yokogawa – Andor), using Zeiss 40X and 63X water objectives.

#### Laser ablation experiments

Laser ablation experiments were performed using a spinning disk microscope equipped with a CSU-XI spinning disk head, using a 63X oil objective. The ablation was done using a pulsed 355 nm ultraviolet laser at a power of 30% and a thickness of 3, interfaced with an iLas system (Roper Scientific) piloted in Metamorph.

#### Two-dimensional models for spindle orientation

2D models to predict spindle orientation were adapted from (Bosveld *et al*., 2016; Minc *et al*., 2011). These models were developed and executed through Matlab (Mathworks) scripts which can be made available upon demand. Starting from the shape of a cell, the model positioned spindle poles and traced MT asters radiating from spindle poles to the contour. For length limitations, we added a fixed maximal length to MTs, normalized to cell length along the A-B axis, which was varied in figure 2e. Each MT is associated to a force, f_MT_, which varies depending on hypothesis. For models based on shape sensing, we posited that f_MT_ =aL_MT_^2^ with L_MT_ the length of the MT and *a*, an arbitrary constant. The scaling to the square was chosen as it best represents a length-dependent system in which MTs pull at the surface (Hara and Kimura, 2009; Minc *et al*., 2011). We note however that other models previously proposed to center and orient spindles with the long axis, including those based on pulling in bulk cytoplasm or MT pushing at the surface and limited by buckling, yielded to the same outputs. For NuMA based models, the force per MT was f_MT_ =b[NuMA]*L_MT_^2^, with b an arbitrary constant, and [NuMA] is the local concentration of NuMA around cells, inputted as a normalized Gaussian distribution with a peak located at the basal pole of cells. The script then looped to compute the torque exert generated by all MTs as a function of all possible spindle angles, and computed a rotational energy potential as a primitive of the torque (Thery *et al*., 2007) (Figure 2f). Free parameters in the model were the number of MTs, the size of the spindle and the spatial extension of asters, which had little influence on model prediction (Figure S3a-d). For the NuMA model, we added one other parameter which is the width of the Gaussian, which was estimated from experimental images, and which did not influence prediction for domain size relatively small as compared to cell contour. Key parameters that altered spindle orientation outputs were thus cell shape, and the asymmetry of spindle positioning, as well as the maximal length of MTs related to cell length (Figure 2g-h and Figure S3a-d)

#### Three-dimensional simulations for spindle position and orientation

The simulation package used for 3D models was developed in Matlab (Mathworks) and was adapted from (Ershov and Minc, 2019; Pierre *et al*., 2016). This package includes a module to extract and reconstitute 3D shapes, spindle pole positions and polarity domains from segmented experimental image, and to add predefined hypothesis similar to those used above in 2D. In 3D, the exclusion of MTs from the basal pole was introduced as a gradient from the basal pole of length limitation, which was scaled to cell length along the A-B axis. The size of spindles and polar basal domains were defined directly from each experimental stack. Once parameters were defined as inputs, the model placed the spindle in a random position and orientation, traced MTs from spindle poles to measure their length and associated a force to each, and computed both global forces and torques exerted by MT asters. To search for minima, the model used a random walk, which follows the minimization of torques and forces. This was achieved by randomly modulating one of the 5 spatial parameters (3 for the position and 2 for the orientation of spindles), and recalculating the force torque at the subsequent step. The simulations followed the direction of force/torque minimization and stopped at a position/orientation once the iteration returns to this equilibrium after a given number of iterations (e.g. 300 runs) (Figure 4e-g, Figure S5a-c, and Videos S4-5). The length of the simulations, the duration needed to identify a stable equilibrium, the noise added to explore other parameters, and other intrinsic parameters of the loop, can be modulated, but were fixed for all the simulations performed in this work. Finally, to ensure that the spindle position/orientation identified did not correspond to a local minimum which could have been biased by the initial conditions, the simulations were run typically 3-4 times from another random starting position/orientation. Finally, once a position was found, the model recalculated the torque landscapes as a function of the 2 possible angles (Figure S5c).

#### Segmentation and 3D rendering

Cell shape or spindle pole 3D segmentations are performed based on confocal z-stacks of cell membranes (i.e. E-cadherin, EpCAM) or MTs (i.e. α-tubulin, NuMA), respectively. Segmentations are performed manually to guarantee the best match with the initial data. Masks corresponding to cell shapes, spindle poles, organoid lumen and basal environment (“exterior”) were generated on each slice of confocal z-stacks using ImageJ.

For generating the 3D rendering of organoid cells, mask outlines were translated into meshes, which are 3D surfaces connecting the perimeters on each slice, by using a Python program adapted from an initial program provided by Emmanuel Faure (LIRMM, Montpellier). Briefly, for a given cell, the mask was first translated into a list of points lying on the perimeters for every position along z where the cell is present. The list of points was then translated into a list of polygonal faces linking the points together. Visualization of meshes was done with the Meshlab software (www.meshlab.net).

Measurements of geometric properties of the segmented cells, in particular their volume V and surface area A, were provided by Meshlab and used to compute the cell sphericity: s=π1/3(6V)2/3/A (Wadell, 1935). From each pixel of the segmented cells, we computed the distance to the lumen. For each cell, the mean cell height H in the apico-basal direction was computed as the difference between the furthest and closest pixels from the lumen, averaged for every position along z where the cell is present. The mean cell width was computed from the cell volume and height assuming a cylindrical geometry and using W=2(V/πH)½. The mean spindle positioning with respect to the apico-basal axis was computed by averaging the distance from each point of the spindle to the lumen.

Multicellular contact distribution was computed from the ImageJ organoid cell shape masks and by using a custom Python program. The multicellular contacts were identified as any pixel in the vicinity of the borders of three or more cell masks (within 20 pixels). The “degree” of the multicellular contact (3-cells, 4-cells etc.) was given by the number of different cell masks in the vicinity of each multicellular contact. Positioning of computed multicellular contacts was then visualized together with cell meshes in Meshlab.

#### Expansion microscopy of organoids

Expansion microscopy protocol was adapted from Gambarotto et al. (Gambarotto et al., 2019). Matrigel domes containing mouse organoids were dissociated by using Gentle Cell Dissociation Reagent. After centrifugation, Matrigel debris were eliminated with the supernatant and organoid pellet kept. Organoids were incubated overnight at 37°C in PBS containing 2% acrylamide / 1.5% formaldehyde.

For gelation, organoids were washed in PBS and pelleted by centrifugation. After resuspension PBS, organoids were layed on a parafilm piece in a humid chamber. After PBS removal with a tip, gelation solution (19.3% sodium acrylate / 10% acrylamide / 0.1% N,N’-methylenbisacrylamide / 0.5% TEMED / 0.1% Ammonium persulfate, in PBS) was added. Gelation solution containing organoids was covered with a 5mm coverslip, incubated 5 min on ice and then 1 hour at 37°C. The coverslip and the gel were placed in a 24-well plate filled with 500μl of denaturation buffer (200mM SDS / 200mM NaCl / 50mM Tris water, PH9). After 15 min agitation at room temperature, the gel detached from the coverslip, was placed in a 1.5ml microtube filled with denaturation buffer and incubated at 95°C for 1h30min.

For immunofluorescence, organoids were transferred in a 24-well plate, blocked and permeabilized during incubation in 1% BSA / 0.025% saponin, in PBS solution for 1 hour at room temperature. Primary antibodies were incubated overnight at 4°C in the 1% BSA / 0.025% saponin, in PBS solution, under gentle shacking. The day after, after 3 PBS washes, secondary antibodies were incubated 4 hours at 4°C and then 1 hour at room temperature upon gentle shacking. After three PBS washes, nuclei were labelled with Hoechst 33342.

For the organoid expansion, the gel was transferred in a 6-well plate filled with ddH2O and wash for 1 hour without agitation. After ddH2O change, the gel was incubated overnight at room temperature. The day after, the expansion index was evaluated based on gel length before and after expansion. For image acquisition, the expanded gel was placed in a poly L-lysine-coated glass chamber after exceeding water removal.

### Statistical analysis

All statistical analyses were performed using Prism (GraphPad Software, San Diego, CA, USA, version 9.0). All circular graphs and statistics were performed using Oriana version 4 (Kovach Computing). Unless otherwise stated, experiments were replicated 3 times independently.

## SUPPLEMENTAL VIDEOS

**Video S1:**

**Description**: Time lapse imaging of α-tubulin-GFP (green) dynamics during mitosis in H2BmCherry (blue) organoids. Images were acquired every 30 sec. Frame rate is 15 fps.

**Video S2:**

**Description**: Time lapse imaging of myosin-IIA-GFP in intestinal organoids. Scale bar, 10μm. Images were acquired every 15 sec. Frame rate is 15 fps.

**Video S3:**

**Description**: Time lapse imaging of EB3-RFP in prophase (left panel) and metaphase (right panel) cells in intestinal organoids. Images were acquired every 4 or 1 sec, respectively. Frame rate is 10 fps.

**Video S4:**

**Description**: Time-lapses of 3D gradient descent simulations to predict planar spindle orientation, started with spindles close to the apical pole (left) or the basal pole (right). Note the similar final predicted equilibrium.

**Video S5:**

**Description**: 3D rotating views of spindle orientations in a representative experimental cell, and as predicted by 3D simulations based on the different indicated hypothesis (NuMA polarity, Full cell shape, or cell shape coupled to length-limitation in MTs).

## REFERENCES

Aigouy, B., Farhadifar, R., Staple, D.B., Sagner, A., Roper, J.C., Julicher, F., and Eaton, S. (2010). Cell flow reorients the axis of planar polarity in the wing epithelium of Drosophila. Cell 142, 773–786. 10.1016/j.cell.2010.07.042.

Artegiani, B., and Clevers, H. (2018). Use and application of 3D-organoid technology. Hum Mol Genet 27, R99–R107. 10.1093/hmg/ddy187.

Bellis, J., Duluc, I., Romagnolo, B., Perret, C., Faux, M.C., Dujardin, D., Formstone, C., Lightowler, S., Ramsay, R.G., Freund, J.N., and De Mey, J.R. (2012). The tumor suppressor Apc controls planar cell polarities central to gut homeostasis. J Cell Biol 198, 331–341. 10.1083/jcb.201204086.

Bergstralh, D.T., and St Johnston, D. (2014). Spindle orientation: what if it goes wrong? Semin Cell Dev Biol 34, 140–145. 10.1016/j.semcdb.2014.06.014.

Biggs, L.C., Kim, C.S., Miroshnikova, Y.A., and Wickstrom, S.A. (2020). Mechanical Forces in the Skin: Roles in Tissue Architecture, Stability, and Function. J Invest Dermatol 140, 284–290. 10.1016/j.jid.2019.06.137.

Bosveld, F., Markova, O., Guirao, B., Martin, C., Wang, Z., Pierre, A., Balakireva, M., Gaugue, I., Ainslie, A., Christophorou, N., et al. (2016). Epithelial tricellular junctions act as interphase cell shape sensors to orient mitosis. Nature 530, 495–498. 10.1038/nature16970.

Campinho, P., Behrndt, M., Ranft, J., Risler, T., Minc, N., and Heisenberg, C.P. (2013). Tension-oriented cell divisions limit anisotropic tissue tension in epithelial spreading during zebrafish epiboly. Nat Cell Biol 15, 1405–1414. 10.1038/ncb2869.

Carminati, M., Gallini, S., Pirovano, L., Alfieri, A., Bisi, S., and Mapelli, M. (2016). Concomitant binding of Afadin to LGN and F-actin directs planar spindle orientation. Nat Struct Mol Biol 23, 155–163. 10.1038/nsmb.3152.

Carroll, T.D., Langlands, A.J., Osborne, J.M., Newton, I.P., Appleton, P.L., and Nathke, I. (2017). Interkinetic nuclear migration and basal tethering facilitates post-mitotic daughter separation in intestinal organoids. J Cell Sci 130, 3862–3877. 10.1242/jcs.211656.

Caussinus, E., and Gonzalez, C. (2005). Induction of tumor growth by altered stem-cell asymmetric division in Drosophila melanogaster. Nat Genet 37, 1125–1129. 10.1038/ng1632.

Chanet, S., Sharan, R., Khan, Z., and Martin, A.C. (2017). Myosin 2-Induced Mitotic Rounding Enables Columnar Epithelial Cells to Interpret Cortical Spindle Positioning Cues. Curr Biol 27, 3350–3358 e3353. 10.1016/j.cub.2017.09.039.

Compton, D.A., and Cleveland, D.W. (1993). NuMA is required for the proper completion of mitosis. J Cell Biol 120, 947–957.

da Silva, S.M., and Vincent, J.P. (2007). Oriented cell divisions in the extending germband of Drosophila. Development 134, 3049–3054. 10.1242/dev.004911.

David, N.B., Martin, C.A., Segalen, M., Rosenfeld, F., Schweisguth, F., and Bellaiche, Y. (2005). Drosophila Ric-8 regulates Galphai cortical localization to promote Galphai-dependent planar orientation of the mitotic spindle during asymmetric cell division. Nat Cell Biol 7, 1083–1090. 10.1038/ncb1319.

di Pietro, F., Echard, A., and Morin, X. (2016). Regulation of mitotic spindle orientation: an integrated view. EMBO Rep 17, 1106–1130. 10.15252/embr.201642292.

Dimitracopoulos, A., Srivastava, P., Chaigne, A., Win, Z., Shlomovitz, R., Lancaster, O.M., Le Berre, M., Piel, M., Franze, K., Salbreux, G., and Baum, B. (2020). Mechanochemical Crosstalk Produces Cell-Intrinsic Patterning of the Cortex to Orient the Mitotic Spindle. Curr Biol 30, 3687–3696 e3684. 10.1016/j.cub.2020.06.098.

Dona, F., Eli, S., and Mapelli, M. (2022). Insights Into Mechanisms of Oriented Division From Studies in 3D Cellular Models. Front Cell Dev Biol 10, 847801. 10.3389/fcell.2022.847801.

Dow, L.E., O’Rourke, K.P., Simon, J., Tschaharganeh, D.F., van Es, J.H., Clevers, H., and Lowe, S.W. (2015). Apc Restoration Promotes Cellular Differentiation and Reestablishes Crypt Homeostasis in Colorectal Cancer. Cell 161, 1539–1552. 10.1016/j.cell.2015.05.033.

Ershov, D., and Minc, N. (2019). Modeling Embryonic Cleavage Patterns. Methods Mol Biol 1920, 393–406. 10.1007/978-1-4939-9009-2_24.

Farhadifar, R., Yu, C.H., Fabig, G., Wu, H.Y., Stein, D.B., Rockman, M., Muller-Reichert, T., Shelley, M.J., and Needleman, D.J. (2020). Stoichiometric interactions explain spindle dynamics and scaling across 100 million years of nematode evolution. Elife 9. 10.7554/eLife.55877.

Farin, H.F., Van Es, J.H., and Clevers, H. (2012). Redundant sources of Wnt regulate intestinal stem cells and promote formation of Paneth cells. Gastroenterology 143, 1518–1529 e1517. 10.1053/j.gastro.2012.08.031.

Fatehullah, A., Tan, S.H., and Barker, N. (2016). Organoids as an in vitro model of human development and disease. Nat Cell Biol 18, 246–254. 10.1038/ncb3312.

Feng, Y., Sentani, K., Wiese, A., Sands, E., Green, M., Bommer, G.T., Cho, K.R., and Fearon, E.R. (2013). Sox9 induction, ectopic Paneth cells, and mitotic spindle axis defects in mouse colon adenomatous epithelium arising from conditional biallelic Apc inactivation. Am J Pathol 183, 493–503. 10.1016/j.ajpath.2013.04.013.

Fleming, E.S., Zajac, M., Moschenross, D.M., Montrose, D.C., Rosenberg, D.W., Cowan, A.E., and Tirnauer, J.S. (2007). Planar spindle orientation and asymmetric cytokinesis in the mouse small intestine. J Histochem Cytochem 55, 1173–1180. 10.1369/jhc.7A7234.2007.

Gambarotto, D., Zwettler, F.U., Le Guennec, M., Schmidt-Cernohorska, M., Fortun, D., Borgers, S., Heine, J., Schloetel, J.G., Reuss, M., Unser, M., et al. (2019). Imaging cellular ultrastructures using expansion microscopy (U-ExM). Nat Methods 16, 71–74. 10.1038/s41592-018-0238-1.

Gehart, H., and Clevers, H. (2019). Tales from the crypt: new insights into intestinal stem cells. Nat Rev Gastroenterol Hepatol 16, 19–34. 10.1038/s41575-018-0081-y.

Gloerich, M., Bianchini, J.M., Siemers, K.A., Cohen, D.J., and Nelson, W.J. (2017). Cell division orientation is coupled to cell-cell adhesion by the E-cadherin/LGN complex. Nat Commun 8, 13996. 10.1038/ncomms13996.

Godard, B.G., Dumollard, R., Heisenberg, C.P., and McDougall, A. (2021). Combined effect of cell geometry and polarity domains determines the orientation of unequal division. Elife 10. 10.7554/eLife.75639.

Godard, B.G., and Heisenberg, C.P. (2019). Cell division and tissue mechanics. Curr Opin Cell Biol 60, 114–120. 10.1016/j.ceb.2019.05.007.

Goldspink, D.A., Matthews, Z.J., Lund, E.K., Wileman, T., and Mogensen, M.M. (2017). Immuno-fluorescent Labeling of Microtubules and Centrosomal Proteins in Ex Vivo Intestinal Tissue and 3D In Vitro Intestinal Organoids. J Vis Exp. 10.3791/56662.

Grill, S.W., and Hyman, A.A. (2005). Spindle positioning by cortical pulling forces. Dev Cell 8, 461–465. 10.1016/j.devcel.2005.03.014.

Hara, Y., and Kimura, A. (2009). Cell-size-dependent spindle elongation in the Caenorhabditis elegans early embryo. Curr Biol 19, 1549–1554. 10.1016/j.cub.2009.07.050.

Hart, K.C., Tan, J., Siemers, K.A., Sim, J.Y., Pruitt, B.L., Nelson, W.J., and Gloerich, M. (2017). E-cadherin and LGN align epithelial cell divisions with tissue tension independently of cell shape. Proc Natl Acad Sci U S A 114, E5845–E5853. 10.1073/pnas.1701703114.

Haupt, A., and Minc, N. (2018). How cells sense their own shape - mechanisms to probe cell geometry and their implications in cellular organization and function. J Cell Sci 131. 10.1242/jcs.214015.

Jacobelli, J., Friedman, R.S., Conti, M.A., Lennon-Dumenil, A.M., Piel, M., Sorensen, C.M., Adelstein, R.S., and Krummel, M.F. (2010). Confinement-optimized three-dimensional T cell amoeboid motility is modulated via myosin IIA-regulated adhesions. Nat Immunol 11, 953–961. 10.1038/ni.1936.

Jimenez, A.J., Schaeffer, A., De Pascalis, C., Letort, G., Vianay, B., Bornens, M., Piel, M., Blanchoin, L., and Thery, M. (2021). Acto-myosin network geometry defines centrosome position. Curr Biol 31, 1206–1220 e1205. 10.1016/j.cub.2021.01.002.

Kelkar, M., Bohec, P., and Charras, G. (2020). Mechanics of the cellular actin cortex: From signalling to shape change. Curr Opin Cell Biol 66, 69–78. 10.1016/j.ceb.2020.05.008.

Kiyomitsu, T., and Cheeseman, I.M. (2012). Chromosome- and spindle-pole-derived signals generate an intrinsic code for spindle position and orientation. Nat Cell Biol 14, 311–317. 10.1038/ncb2440.

Kotak, S., Busso, C., and Gonczy, P. (2012). Cortical dynein is critical for proper spindle positioning in human cells. J Cell Biol 199, 97–110. 10.1083/jcb.201203166.

Krndija, D., El Marjou, F., Guirao, B., Richon, S., Leroy, O., Bellaiche, Y., Hannezo, E., and Matic Vignjevic, D. (2019). Active cell migration is critical for steady-state epithelial turnover in the gut. Science 365, 705–710. 10.1126/science.aau3429.

Lancaster, M.A., and Knoblich, J.A. (2012). Spindle orientation in mammalian cerebral cortical development. Curr Opin Neurobiol 22, 737–746. 10.1016/j.conb.2012.04.003.

Lancaster, O.M., Le Berre, M., Dimitracopoulos, A., Bonazzi, D., Zlotek-Zlotkiewicz, E., Picone, R., Duke, T., Piel, M., and Baum, B. (2013). Mitotic rounding alters cell geometry to ensure efficient bipolar spindle formation. Dev Cell 25, 270–283. 10.1016/j.devcel.2013.03.014.

Lechler, T., and Fuchs, E. (2005). Asymmetric cell divisions promote stratification and differentiation of mammalian skin. Nature 437, 275–280. 10.1038/nature03922.

Lechler, T., and Mapelli, M. (2021). Spindle positioning and its impact on vertebrate tissue architecture and cell fate. Nat Rev Mol Cell Biol 22, 691–708. 10.1038/s41580-021-00384-4.

Lough, K.J., Byrd, K.M., Descovich, C.P., Spitzer, D.C., Bergman, A.J., Beaudoin, G.M., 3rd, Reichardt, L.F., and Williams, S.E. (2019). Telophase correction refines division orientation in stratified epithelia. Elife 8. 10.7554/eLife.49249.

Luxenburg, C., Pasolli, H.A., Williams, S.E., and Fuchs, E. (2011). Developmental roles for Srf, cortical cytoskeleton and cell shape in epidermal spindle orientation. Nat Cell Biol 13, 203–214. 10.1038/ncb2163.

Macara, I.G., Guyer, R., Richardson, G., Huo, Y., and Ahmed, S.M. (2014). Epithelial homeostasis. Curr Biol 24, R815–825. 10.1016/j.cub.2014.06.068.

Mao, Y., Tournier, A.L., Hoppe, A., Kester, L., Thompson, B.J., and Tapon, N. (2013). Differential proliferation rates generate patterns of mechanical tension that orient tissue growth. EMBO J 32, 2790–2803. 10.1038/emboj.2013.197.

McCaffrey, L.M., and Macara, I.G. (2011). Epithelial organization, cell polarity and tumorigenesis. Trends Cell Biol 21, 727–735. 10.1016/j.tcb.2011.06.005.

McHugh, T., Gluszek, A.A., and Welburn, J.P.I. (2018). Microtubule end tethering of a processive kinesin-8 motor Kif18b is required for spindle positioning. J Cell Biol 217, 2403–2416. 10.1083/jcb.201705209.

McKinley, K.L., Stuurman, N., Royer, L.A., Schartner, C., Castillo-Azofeifa, D., Delling, M., Klein, O.D., and Vale, R.D. (2018). Cellular aspect ratio and cell division mechanics underlie the patterning of cell progeny in diverse mammalian epithelia. Elife 7. 10.7554/eLife.36739.

McNally, F.J. (2013). Mechanisms of spindle positioning. J Cell Biol 200, 131–140. 10.1083/jcb.201210007.

Minc, N., Burgess, D., and Chang, F. (2011). Influence of cell geometry on division-plane positioning. Cell 144, 414–426. 10.1016/j.cell.2011.01.016.

Mitchison, T.J., Ishihara, K., Nguyen, P., and Wuhr, M. (2015). Size Scaling of Microtubule Assemblies in Early Xenopus Embryos. Cold Spring Harb Perspect Biol 7, a019182. 10.1101/cshperspect.a019182.

Morin, X., and Bellaiche, Y. (2011). Mitotic spindle orientation in asymmetric and symmetric cell divisions during animal development. Dev Cell 21, 102–119. 10.1016/j.devcel.2011.06.012.

Morin, X., Jaouen, F., and Durbec, P. (2007). Control of planar divisions by the G-protein regulator LGN maintains progenitors in the chick neuroepithelium. Nat Neurosci 10, 1440–1448. 10.1038/nn1984.

Muzumdar, M.D., Tasic, B., Miyamichi, K., Li, L., and Luo, L. (2007). A global double-fluorescent Cre reporter mouse. Genesis 45, 593–605. 10.1002/dvg.20335.

Nakajima, Y., Meyer, E.J., Kroesen, A., McKinney, S.A., and Gibson, M.C. (2013). Epithelial junctions maintain tissue architecture by directing planar spindle orientation. Nature 500, 359–362. 10.1038/nature12335.

Niwayama, R., Moghe, P., Liu, Y.J., Fabreges, D., Buchholz, F., Piel, M., and Hiiragi, T. (2019). A Tug-of-War between Cell Shape and Polarity Controls Division Orientation to Ensure Robust Patterning in the Mouse Blastocyst. Dev Cell 51, 564–574 e566. 10.1016/j.devcel.2019.10.012.

Norden, C., Young, S., Link, B.A., and Harris, W.A. (2009). Actomyosin is the main driver of interkinetic nuclear migration in the retina. Cell 138, 1195–1208. 10.1016/j.cell.2009.06.032.

O., H. (1884). Das Problem der Befruchtung und der Isotropie des Eies, eine Theory der Vererbung. Jenaische Zeitschrift fuer Naturwissenschaft 18, 21–23.

Peyre, E., Jaouen, F., Saadaoui, M., Haren, L., Merdes, A., Durbec, P., and Morin, X. (2011). A lateral belt of cortical LGN and NuMA guides mitotic spindle movements and planar division in neuroepithelial cells. J Cell Biol 193, 141–154. 10.1083/jcb.201101039.

Pierre, A., Salle, J., Wuhr, M., and Minc, N. (2016). Generic Theoretical Models to Predict Division Patterns of Cleaving Embryos. Dev Cell 39, 667–682. 10.1016/j.devcel.2016.11.018.

Quyn, A.J., Appleton, P.L., Carey, F.A., Steele, R.J., Barker, N., Clevers, H., Ridgway, R.A., Sansom, O.J., and Nathke, I.S. (2010). Spindle orientation bias in gut epithelial stem cell compartments is lost in precancerous tissue. Cell Stem Cell 6, 175–181. 10.1016/j.stem.2009.12.007.

Salle, J., and Minc, N. (2021). Cell division geometries as central organizers of early embryo development. Semin Cell Dev Biol. 10.1016/j.semcdb.2021.08.004.

Salomon, J., Gaston, C., Magescas, J., Duvauchelle, B., Canioni, D., Sengmanivong, L., Mayeux, A., Michaux, G., Campeotto, F., Lemale, J., et al. (2017). Contractile forces at tricellular contacts modulate epithelial organization and monolayer integrity. Nat Commun 8, 13998. 10.1038/ncomms13998.

Sato, T., Stange, D.E., Ferrante, M., Vries, R.G., Van Es, J.H., Van den Brink, S., Van Houdt, W.J., Pronk, A., Van Gorp, J., Siersema, P.D., and Clevers, H. (2011a). Long-term expansion of epithelial organoids from human colon, adenoma, adenocarcinoma, and Barrett’s epithelium. Gastroenterology 141, 1762–1772. 10.1053/j.gastro.2011.07.050.

Sato, T., van Es, J.H., Snippert, H.J., Stange, D.E., Vries, R.G., van den Born, M., Barker, N., Shroyer, N.F., van de Wetering, M., and Clevers, H. (2011b). Paneth cells constitute the niche for Lgr5 stem cells in intestinal crypts. Nature 469, 415–418. 10.1038/nature09637.

Scarpa, E., Finet, C., Blanchard, G.B., and Sanson, B. (2018). Actomyosin-Driven Tension at Compartmental Boundaries Orients Cell Division Independently of Cell Geometry In Vivo. Dev Cell 47, 727–740 e726. 10.1016/j.devcel.2018.10.029.

Schaefer, M., Shevchenko, A., Shevchenko, A., and Knoblich, J.A. (2000). A protein complex containing Inscuteable and the Galpha-binding protein Pins orients asymmetric cell divisions in Drosophila. Curr Biol 10, 353–362. 10.1016/s0960-9822(00)00401-2.

Schenk, J., Wilsch-Brauninger, M., Calegari, F., and Huttner, W.B. (2009). Myosin II is required for interkinetic nuclear migration of neural progenitors. Proc Natl Acad Sci U S A 106, 16487–16492. 10.1073/pnas.0908928106.

Seldin, L., and Macara, I. (2017). Epithelial spindle orientation diversities and uncertainties: recent developments and lingering questions. F1000Res 6, 984. 10.12688/f1000research.11370.1.

Spear, P.C., and Erickson, C.A. (2012). Apical movement during interkinetic nuclear migration is a two-step process. Dev Biol 370, 33–41. 10.1016/j.ydbio.2012.06.031.

Stout, J.R., Yount, A.L., Powers, J.A., Leblanc, C., Ems-McClung, S.C., and Walczak, C.E. (2011). Kif18B interacts with EB1 and controls astral microtubule length during mitosis. Mol Biol Cell 22, 3070–3080. 10.1091/mbc.E11-04-0363.

Tan, D.W., and Barker, N. (2014). Intestinal stem cells and their defining niche. Curr Top Dev Biol 107, 77–107. 10.1016/B978-0-12-416022-4.00003-2.

Thery, M., Jimenez-Dalmaroni, A., Racine, V., Bornens, M., and Julicher, F. (2007). Experimental and theoretical study of mitotic spindle orientation. Nature 447, 493–496. 10.1038/nature05786.

Thery, M., Racine, V., Pepin, A., Piel, M., Chen, Y., Sibarita, J.B., and Bornens, M. (2005). The extracellular matrix guides the orientation of the cell division axis. Nat Cell Biol 7, 947–953. 10.1038/ncb1307.

Van Lidth de Jeude, J.F., Vermeulen, J.L., Montenegro-Miranda, P.S., Van den Brink, G.R., and Heijmans, J. (2015). A protocol for lentiviral transduction and downstream analysis of intestinal organoids. J Vis Exp. 10.3791/52531.

Van Ness, J., and Pettijohn, D.E. (1983). Specific attachment of nuclear-mitotic apparatus protein to metaphase chromosomes and mitotic spindle poles: possible function in nuclear reassembly. J Mol Biol 171, 175–205. 10.1016/s0022-2836(83)80352-0.

Verde, F., Dogterom, M., Stelzer, E., Karsenti, E., and Leibler, S. (1992). Control of microtubule dynamics and length by cyclin A- and cyclin B-dependent kinases in Xenopus egg extracts. J Cell Biol 118, 1097–1108. 10.1083/jcb.118.5.1097.

Wadell, H. (1935). Volume, Shape, and Roundness of Quartz Particles. The Journal of Geology 43.

Wee, B., Johnston, C.A., Prehoda, K.E., and Doe, C.Q. (2011). Canoe binds RanGTP to promote Pins(TPR)/Mud-mediated spindle orientation. J Cell Biol 195, 369–376. 10.1083/jcb.201102130.

Wuhr, M., Chen, Y., Dumont, S., Groen, A.C., Needleman, D.J., Salic, A., and Mitchison, T.J. (2008). Evidence for an upper limit to mitotic spindle length. Curr Biol 18, 1256–1261. 10.1016/j.cub.2008.07.092.

Wyatt, T.P., Harris, A.R., Lam, M., Cheng, Q., Bellis, J., Dimitracopoulos, A., Kabla, A.J., Charras, G.T., and Baum, B. (2015). Emergence of homeostatic epithelial packing and stress dissipation through divisions oriented along the long cell axis. Proc Natl Acad Sci U S A 112, 5726–5731. 10.1073/pnas.1420585112.

Xiong, F., Ma, W., Hiscock, T.W., Mosaliganti, K.R., Tentner, A.R., Brakke, K.A., Rannou, N., Gelas, A., Souhait, L., Swinburne, I.A., et al. (2014). Interplay of cell shape and division orientation promotes robust morphogenesis of developing epithelia. Cell 159, 415–427. 10.1016/j.cell.2014.09.007.

Yu, F., Morin, X., Cai, Y., Yang, X., and Chia, W. (2000). Analysis of partner of inscuteable, a novel player of Drosophila asymmetric divisions, reveals two distinct steps in inscuteable apical localization. Cell 100, 399–409. 10.1016/s0092-8674(00)80676-5.

Zhang, Y., Conti, M.A., Malide, D., Dong, F., Wang, A., Shmist, Y.A., Liu, C., Zerfas, P., Daniels, M.P., Chan, C.C., et al. (2012). Mouse models of MYH9-related disease: mutations in nonmuscle myosin II-A. Blood 119, 238–250. 10.1182/blood-2011-06-358853.

Zheng, Z., Zhu, H., Wan, Q., Liu, J., Xiao, Z., Siderovski, D.P., and Du, Q. (2010). LGN regulates mitotic spindle orientation during epithelial morphogenesis. J Cell Biol 189, 275–288. 10.1083/jcb.200910021.

Zhu, J., Burakov, A., Rodionov, V., and Mogilner, A. (2010). Finding the cell center by a balance of dynein and myosin pulling and microtubule pushing: a computational study. Mol Biol Cell 21, 4418–4427. 10.1091/mbc.E10-07-0627.

