## Supplementary information for "Length-limitation of astral microtubules orients cell divisions in intestinal crypts"

### SUPPLEMENTAL INFORMATION

#### Supplementary Figures

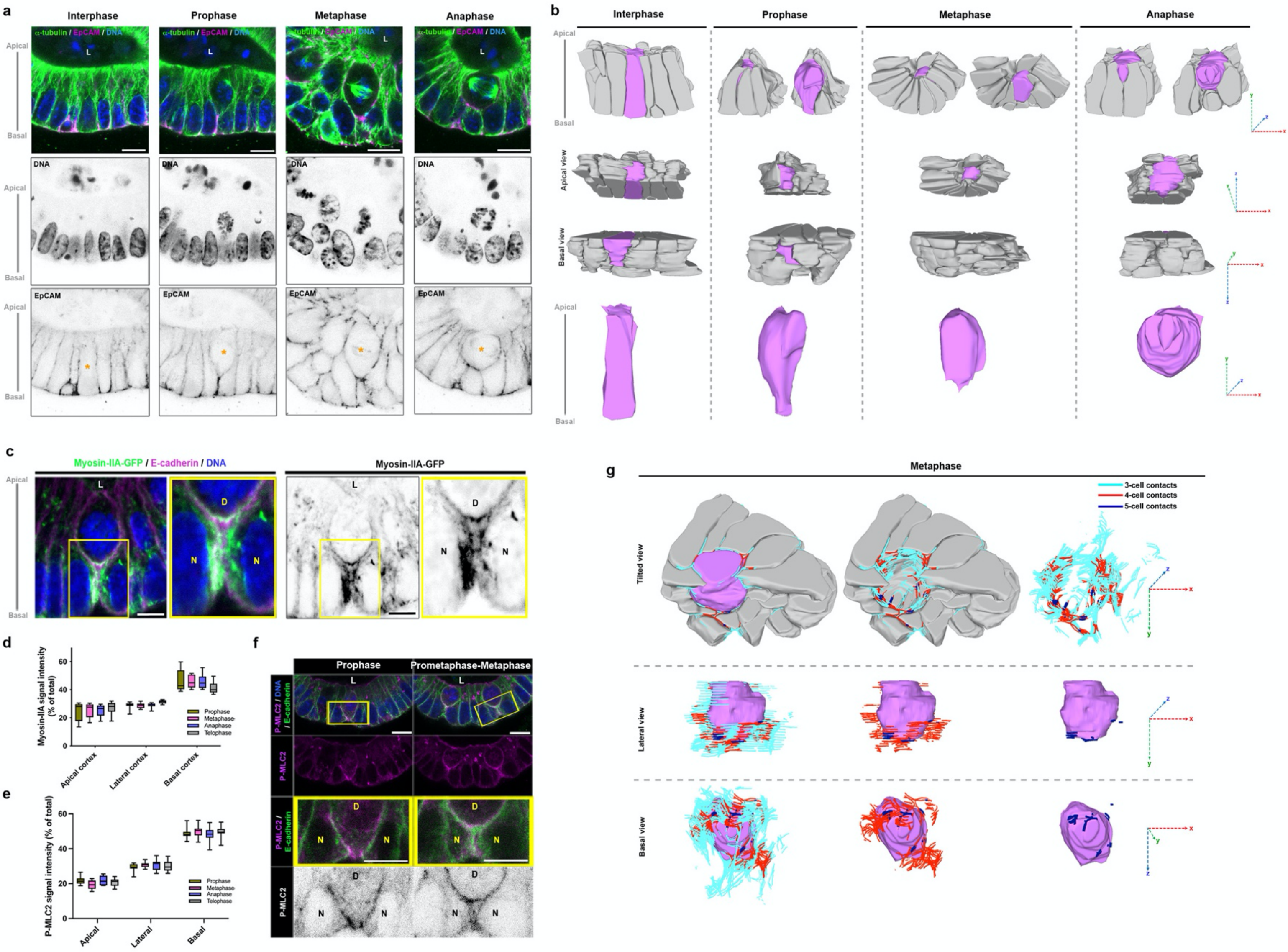


Saleh et al., Figure S1

**Figure S1. Cell rearrangements and mitotic cell shape changes during mitosis in organoid crypt-like structures. a** Confocal analysis of α-tubulin (green) and EpCAM (magenta) in interphase, prophase, metaphase or anaphase cells in intestinal organoids. Nuclei are stained in blue. Orange asterisk pinpoints the cells of interest. L, lumen. Scale bars, 5 μm. **b** Representative 3D rendering of dividing cells (magenta) and neighboring cells (grey) after segmentation of cell membranes based on confocal z-stacks in crypt organoids. Are shown apico-basal, apical or basal views. Spatial coordinates are shown. **c** Confocal analysis of myosin-IIA-KI-GFP (green) and E-cadherin (magenta) in a metaphase cell. L, lumen. D, dividing cell. N, neighboring cell. Scale bar, 5μm. **d** Analysis of the percentage of total signal intensity for myosin-IIA-KI-GFP at the apical, lateral or basal cortex during mitotic steps. n = 10 cells. **e** Analysis of the percentage of total signal intensity of P-MLC2 in the apical, lateral or basal cortex of prophase, metaphase, anaphase and telophase cells. n = 16 prophase cells, 14 metaphase cells, 11 anaphase cells, and 11 telophase cells. **f** Confocal analysis of P-MLC2 (magenta) and E-cadherin (green) distribution in prophase and metaphase cells in intestinal organoids. Scale bars, 10μm; scale bars in the inserts, 5μm. **g** Representative 3D rendering of the metaphase cell of interest (magenta) and neighboring cells (grey) after segmentation of cell membranes based on a representative confocal z-stack in intestinal organoids. 3-cell (cyan), 4-cell (red) and 5-cell (blue) contacts are depicted on the tilted, lateral and basal views.

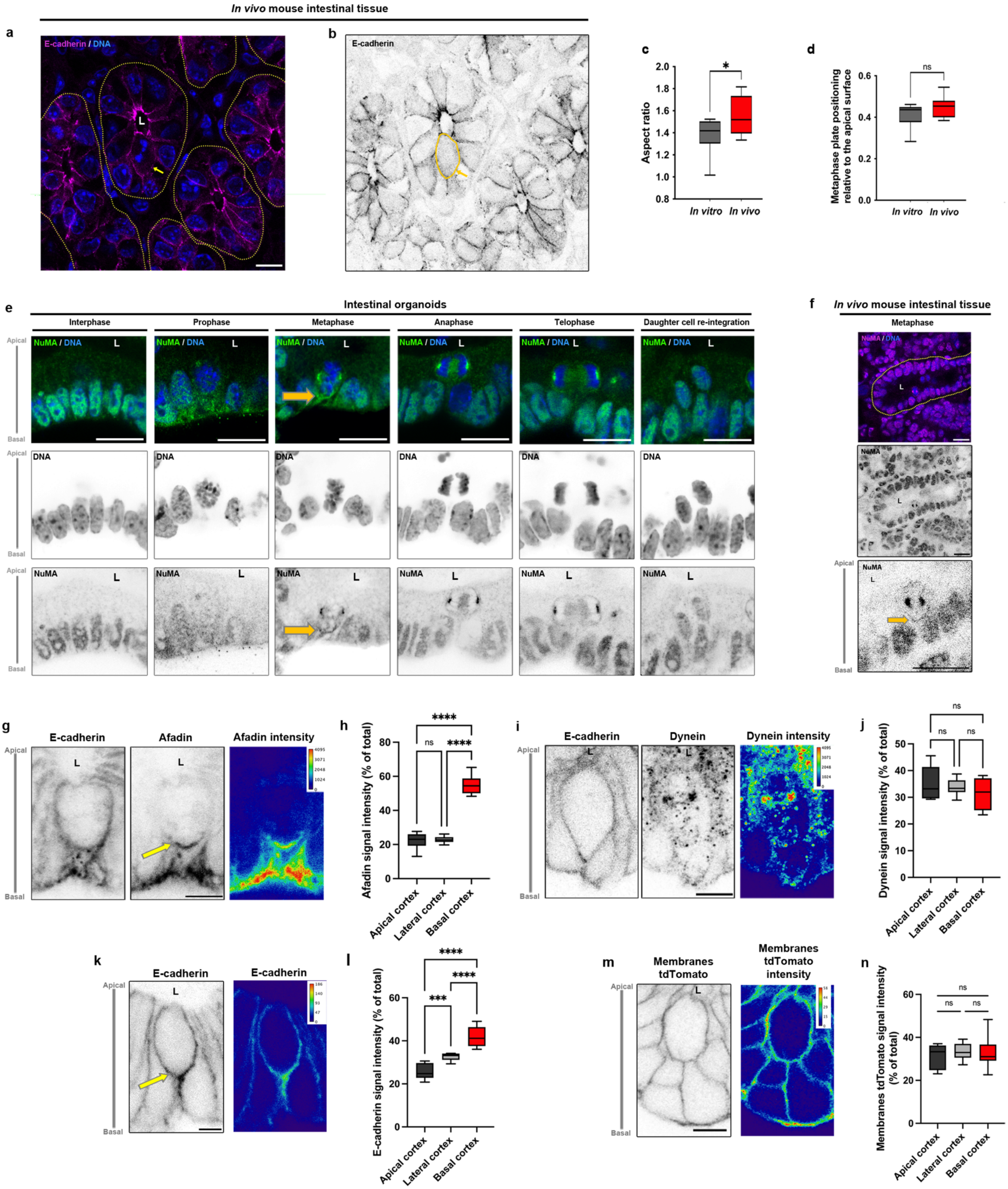


Saleh et al., Figure S2

**Figure S2. Spindle positioning and localization of polarity cues in metaphase cells in mouse jejunum and intestinal organoids. a-b** Confocal analysis of E-cadherin (magenta) localization in mouse intestinal crypts. Nuclei are stained in blue. Yellow dotted lines in **a** delimit crypt compartments. Yellow arrow shows a metaphase cell. Yellow line in **b** surrounds a metaphase cell. Scale bar, 10µm. **c** Statistical analysis of the aspect ratio (distance between the spindle axis and the basal membrane over the distance between the spindle axis and the apical membrane) (n = 10 cells). Aspect ratio (*in vitro*) = 1.378±0.05 (mean±S.E.M), aspect ratio (*in vivo*) = 1.553±0.057. Unpaired t-test, *p = 0.03. **d** Statistical analysis of the metaphase plate positioning (distance between the middle of the metaphase plate and the apical membrane) (n = 10 cells). Metaphase plate positioning (*in vitro*) = 0.411±0.019 (mean±S.E.M), (*in vivo*) = 0.451±0.015. Unpaired t-test, ns, non-significant. **e** Confocal analysis of the distribution of endogenous NuMA (green) during a mitotic cycle in intestinal organoids. Nuclei are stained in blue. Yellow arrow points to NuMA cortical distribution. L, lumen. Scale bar, 15µm. **f** Confocal analysis of the distribution of endogenous NuMA *in vivo* in mouse jejunum crypts. Crypt tissue is delimited by a yellow dotted line. Nuclei are stained in blue. Yellow arrow points to NuMA cortical distribution. L, lumen. Scale bar, 20µm. **g** Confocal analysis of afadin together with E-cadherin in organoid metaphase cells. Afadin signal intensity is color-coded with Fire LUT table from ImageJ on the right panel. Color scale bar indicates the gray value intensity. Yellow arrow points to basal accumulation of afadin. Scale bar, 5 µm. **h** Statistical analysis of afadin signal intensity at the apical, lateral or basal cortex. Signal intensity at the apical cortex = 22.19±1.38 (mean±S.E.M), lateral cortex = 22.94±0.59, basal cortex = 54.83±1.74. n = 10 cells. One-way ANOVA test and Tukey's multiple comparison test, ****p <0.0001. **i** Confocal analysis of dynein together with E-cadherin in organoid metaphase cells. Dynein signal intensity is color-coded with Fire LUT table from ImageJ on the right panel. Color scale bar indicates the gray value intensity. Scale bar, 5 µm. **j** Statistical analysis of dynein signal intensity at the apical, lateral or basal cortex. Signal intensity at the apical cortex = 35.32±2.06 (mean±S.E.M), lateral cortex = 33.72±0.9, basal cortex = 30.94±1.85. n = 10 cells. Two-way ANOVA test and Tukey's multiple comparison test, ns non-significant. **k** Confocal analysis of E-cadherin in organoid metaphase cells. E-cadherin signal intensity is color-coded with Fire LUT table from ImageJ on the right panel. Color scale bar indicates the gray value intensity. Yellow arrow points to basal accumulation of E-cadherin. Scale bar, 5 µm. **l** Statistical analysis of E-cadherin signal intensity at the apical, lateral or basal cortex. Signal intensity at the apical cortex = 25.77±1.15 (mean±S.E.M), lateral cortex = 32.5±0.56, basal cortex = 41.72±1.47. n = 10 cells. Two-way ANOVA test and Tukey's multiple comparison test, ***p = 0.0007, ****p <0.0001. **m** Confocal analysis of Membranes-tdTomato in organoid metaphase cells. tdTomato signal intensity is color-coded with Fire LUT table from ImageJ on the right panel in **c**. Color scale bar indicates the gray value intensity. Scale bar, 5 µm. **n** Statistical analysis of tdTomato signal intensity at the apical, lateral or basal cortex. Signal intensity at the apical cortex = 31.48±1.71 (mean±S.E.M), lateral cortex = 33.27±1.24, basal cortex = 32.84±2.19. n = 10 cells. Two-way ANOVA test and Tukey's multiple comparison test, ns non-significant. For each experiment, three independent experiments were carried out.

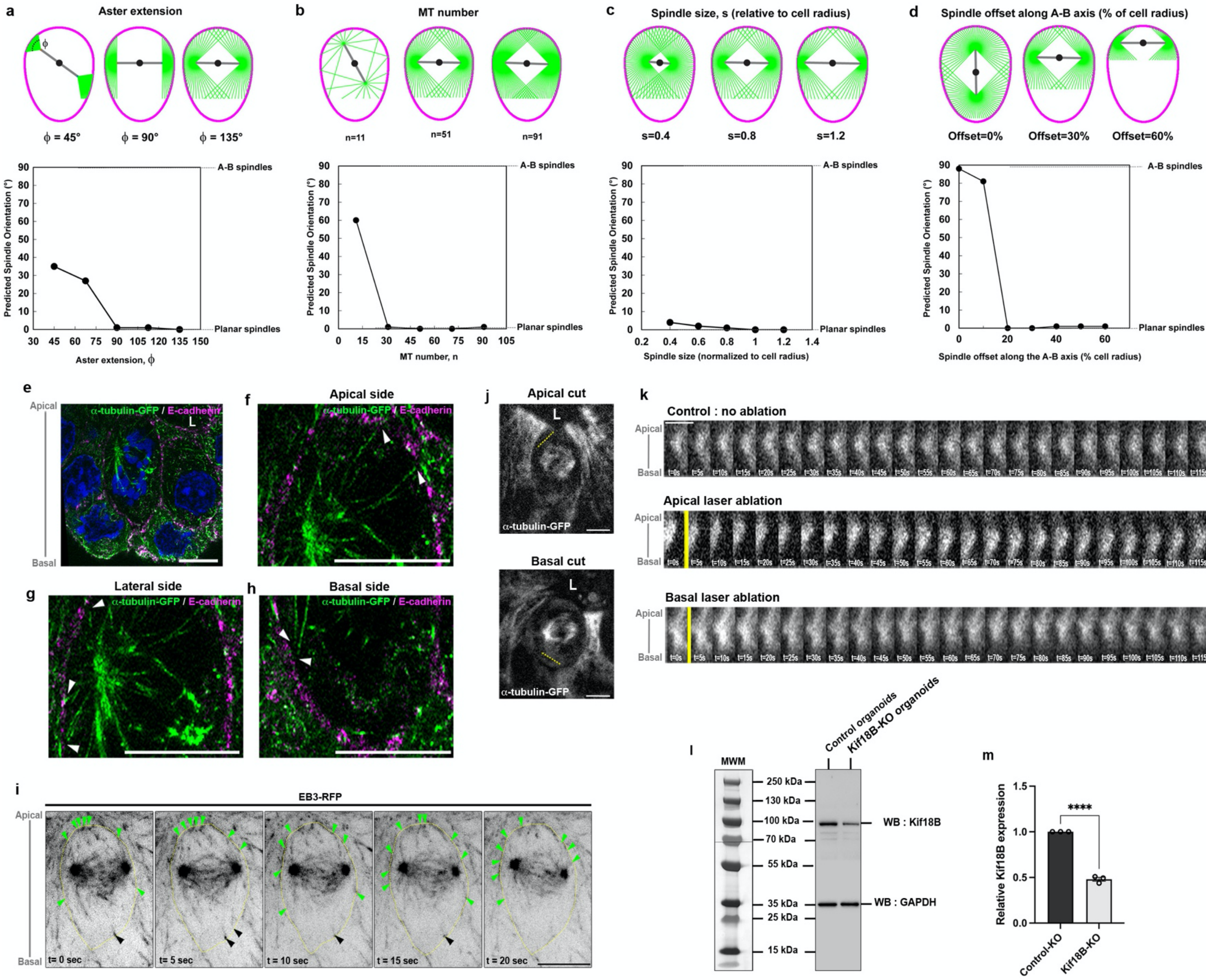


Saleh et al., Figure S3

**Figure S3. Astral MT participation in spindle positioning in intestinal organoids. a** Evolution of the predicted preferred spindle orientation with respect to the tissue plane as a function of aster extension parameter Φ, in the 2D model. **b** Predicted preferred spindle orientation with respect to the tissue plane as a function of MT numbers per aster, n, in the 2D model. **c** Predicted preferred spindle orientation with respect to the tissue plane as a function of spindle size related to cell radii, s, in the 2D model. **d** Predicted preferred spindle orientation with respect to the tissue plane as a function of spindle positional offset along the A-B axis in the 2D model. **e-h** Expansion microscopy and confocal analysis of α-tubulin (green) and E-cadherin (magenta) in organoid metaphase cells. Nuclei (blue) are labeled with Hoechst 33342. Apical (**f**), lateral (**g**) and basal (**h**) cortical domains are presented. Nuclei (blue) are labeled with Hoechst 33342. Corrected scale bar, 5µm. **i** Time-lapse imaging of astral MTs during metaphase in EB3-RFP organoids. Green arrowheads point on astral MTs contacting the apical and lateral cortex, black arrowheads on astral MTs contacting the basal cortex. Metaphase cell shape is delimited by a yellow dotted line. Scale bar, 10µm. **j** Representative spinning disk image of α-tubulin-GFP signal in organoid metaphase cells. Positions of apical or basal laser cut are shown in yellow dotted lines. Scale bar, 2µm. **k** Kymograph analysis of the α-tubulin-GFP signal at the spindle poles in control (no laser ablation), or after apical or basal laser cut in organoid metaphase cells. Yellow vertical line indicates the time of the laser cut. Scale bar, 1µm**. l** Western blot analysis of Kif18B expression in control-KO or Kif18B-KO organoids 3-days after induction. GAPDH was used as a loading control. **m** Statistical analysis of Kif18B expression in control-KO or Kif18B-KO organoids 3-days after induction. Relative expression of Kif18B in Kif18B-KO organoids = 0.479±0.021 (mean±S.E.M). Three independent experiments were carried out. Unpaired t-test, ****p <0.0001.

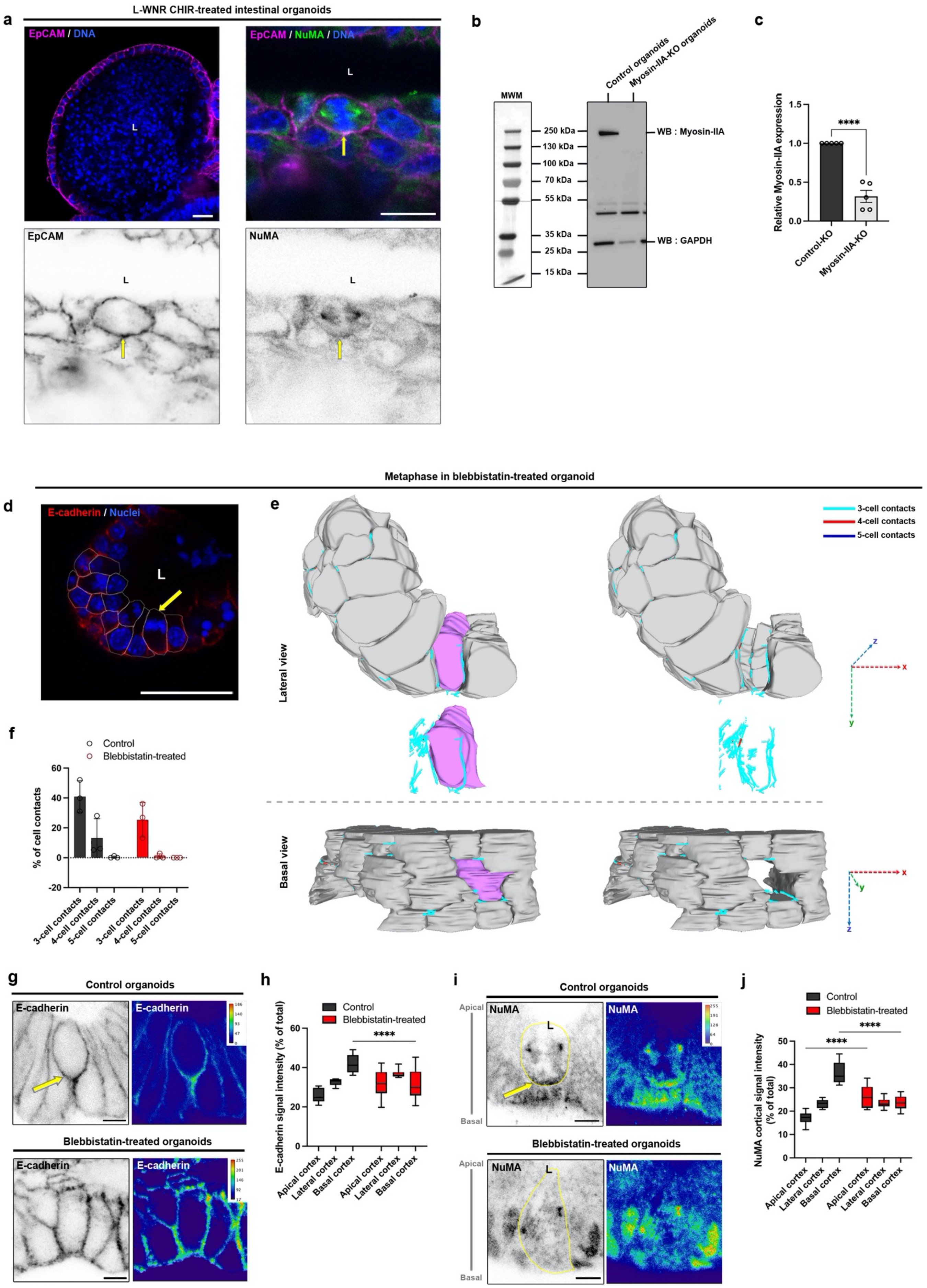


Saleh et al., Figure S4

**Figure S4. Impact of the metaphase cell shape in spindle orientation in intestinal organoids. a** Confocal analysis of EpCAM (magenta) and NuMA (green) distribution in metaphase cells in cystic organoids after L-WNR culture and CHIR treatment. Nuclei are shown in blue. Yellow arrows point to a metaphase cell. Scale bar, 10μm. **b** Western blot analysis of myosin-IIA expression in control-KO or myosin-IIA-KO organoids 3-days after induction. GAPDH was used as a loading control. **c** Statistical analysis of myosin-IIA expression in control-KO or myosin-IIA-KO organoids 3-days after induction. Relative expression of myosin-IIA in myosin-IIA-KO organoids = 0.317±0.077 (mean±S.E.M). Five independent experiments were carried out. Unpaired t-test, ****p <0.0001. **d** Representative confocal image of E-cadherin (red) and nuclei (blue) localization in a crypt organoid after blebbistatin treatment. White lines delimit the cell to be segmented. Yellow arrow shows the metaphase cell of interest. L, lumen. Scale bar, 20 μm. **e** Representative 3D rendering of the metaphase cell of interest shown in **d** (magenta) and neighboring cells (grey) after segmentation of cell membranes based on the representative confocal z-stack in blebbistatin-treated organoids. 3-cell (cyan), 4-cell (red) and 5-cell (blue) contacts are depicted on the lateral and basal views. Spatial coordinates are shown. **f** Quantitative analysis of the proportion of 3-, 4- and 5-cell contacts in control (DMSO-treated) or blebbistatin-treated organoids. **g** Confocal analysis of E-cadherin distribution in metaphase cells in DMSO- or blebbistatin-treated organoids. E-cadherin intensity map was generated with Physics LUT table from ImageJ; color scale bar indicates the gray value intensity. Yellow arrow points to metaphase cell. Scale bar, 5μm**. h** Statistical analysis of E-cadherin distribution in metaphase cells of DMSO- or blebbistatin-treated organoids. Two-way ANOVA and multiple comparisons. ****p <0.0001. **i** Confocal analysis of NuMA distribution in metaphase cells in DMSO- or blebbistatin-treated organoids. NuMA intensity map was generated with Physics LUT table from ImageJ; color scale bar indicates the gray value intensity. Metaphase cell is delimited with a yellow line. Yellow arrowheads point to an increase of NuMA signal at the basal cortex. Scale bar, 5μm**. j** Statistical analysis of NuMA distribution in metaphase cells of DMSO- or blebbistatin-treated organoids. Two-way ANOVA and multiple comparisons. ****p <0.0001. ns, non-significant.

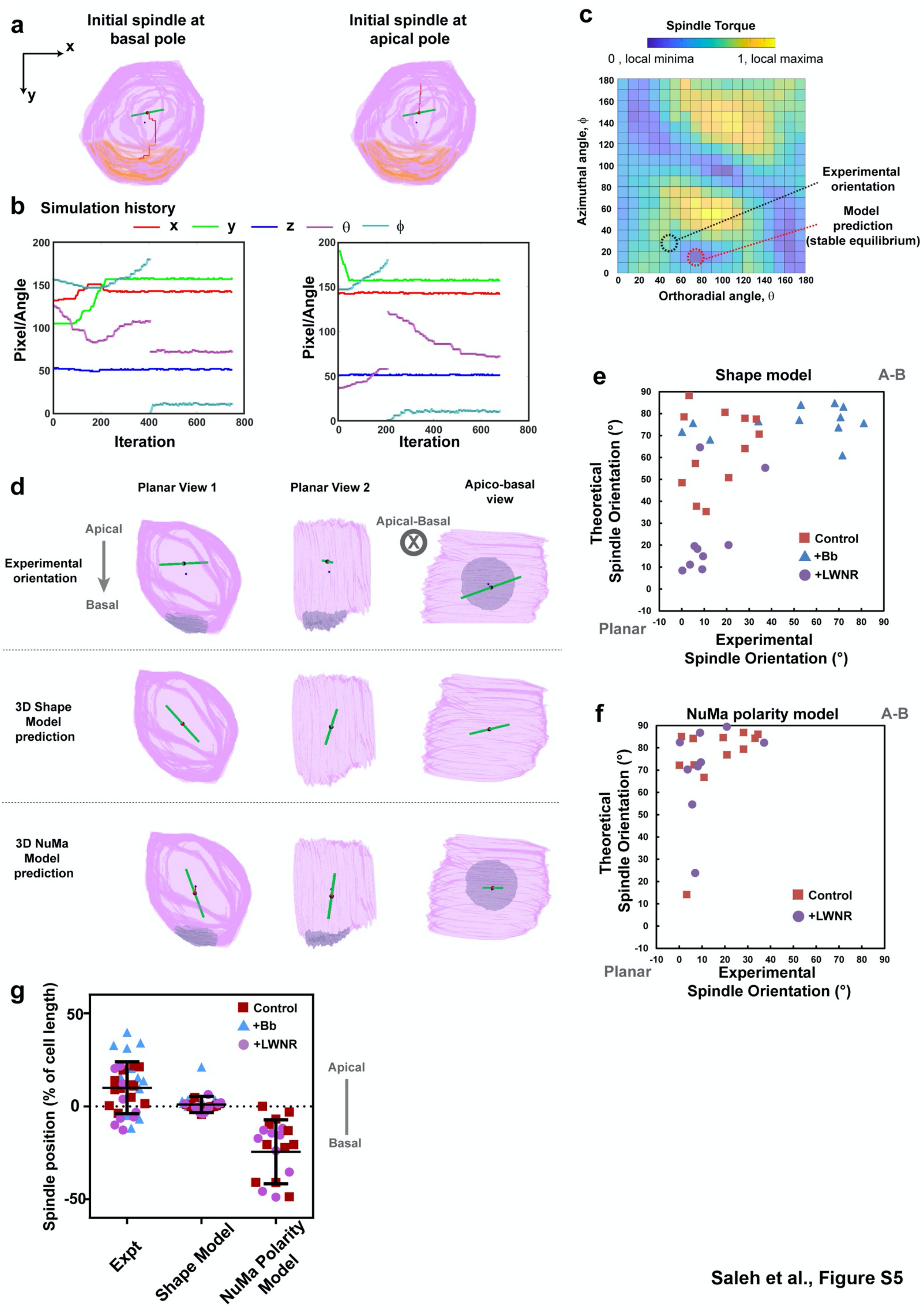


Saleh et al., Figure S5

**Figure S5. 3D simulation of spindle positioning in organoids. a**. 3D simulation for the same cell as presented in Figure 4e, but with initial positions of spindles close to the basal (left) or apical pole (right). The red trace shows the history of the spindle center during the simulations. **b.** Corresponding traces of spindle position (x,y,z) and orientations (θ and Φ) during the gradient descent simulation. Note how the traces become flat at the end of simulations indicating that an equilibrium has been found. **c**. Spindle torque color maps plotted as a function of all possible orthoradial and azimuthal angles, highlighting the identified global minimum and the experimental spindle orientation in 3D. **d.** Experimental and model predicted spindle orientation for the same cell as in Figure 4e, using models based on the full cell shape or based on NuMA polarity at the basal pole. **e.** Predicted spindle orientation angle with respect to the A-B axis, plotted as a function of the experimental axis for 10 individual control, blebbistatin-treated or L-WNR treated crypt cells, for a 3D model based on cell shape. Note the poor agreement for control cells. **f.** Predicted spindle orientation angle with respect to the A-B axis, plotted as a function of the experimental axis for 10 individual control, and L-WNR treated crypt cells, for a 3D model based on cell polarity. Blebbistatin treated cells are absent from this analysis as they fail to properly assemble a basal NuMA domain. Note the poor agreement for both control and L-WNR cells. **g.** Experimental and theoretical prediction of spindle asymmetric position towards the apical cell poles in experiments vs models based on cell shape or NuMA polarity domains.
